# A century of soybean breeding increased photosynthetic capacity but not NPQ relaxation

**DOI:** 10.64898/2026.08.28.747836

**Authors:** Lauana P. de Oliveira, Komal Attri, Lynn Doran, Laurie B. Leonelli, Stephen P. Long, Elizabeth A. Ainsworth

**Affiliations:** Carl R. Woese Institute for Genomic Biology, University of Illinois Urbana-Champaign, Urbana, IL 61801, USA; Department of Agricultural and Biological Engineering, University of Illinois Urbana-Champaign, Urbana, IL 61801, USA; Department of Plant Biology, University of Illinois Urbana-Champaign, Urbana, IL 61801, USA; Department of Crop Sciences, University of Illinois Urbana-Champaign, Urbana, IL 61801, USA

**Keywords:** Photoprotection recovery, Genetic gain, Fluctuating light, Xanthophyll cycle, Photosynthetic efficiency, *Glycine max*

## Abstract

Accelerating photoprotective regulation to improve carbon assimilation is a promising strategy to increase crop productivity. Although rapid non-photochemical quenching (NPQ) relaxation has been validated as a target through metabolic engineering, it remains unclear whether conventional breeding has improved this trait. Here, we investigated whether more than a century of soybean breeding enhanced NPQ relaxation alongside light-saturated carbon assimilation and seed traits. We evaluated a historical panel of 24 soybean genotypes across vegetative and reproductive developmental stages by integrating NPQ relaxation, gas exchange parameters, xanthophyll-cycle pigment profiles, expression of key photoprotective genes (*VDE*, *PsbS*, and *ZEP*), seed number and seed weight. NPQ relaxation parameters were not consistently associated with genotype release year, seed number, or seed weight at either developmental stage. The only exception was the amplitude of the rapidly relaxing NPQ component (AqE), which was negatively correlated with all three variables during the reproductive stage. In contrast, genotype release year was positively associated with maximum net CO₂ assimilation rate (*A*_max_), maximum carboxylation rate of Rubisco (*V*_cmax_), maximum electron transport rate (*J*_max_), seed number, and seed weight, while *A*_max_ and *V*_cmax_ were positively correlated with seed number and seed weight. These findings indicate that the greater photosynthetic capacity of modern genotypes was not accompanied by faster photoprotective response. Thus, photoprotective regulation has not kept pace with gains in photosynthetic capacity under field conditions. We conclude that rapid NPQ relaxation remains an important target for synchronizing photoprotection with the high photosynthetic capacity of modern soybean lines.

## Introduction

Improving photosynthetic efficiency is a key target of biotechnological research aimed to improve global crop yields (Long *et al*., 2025). However, reaching the theoretical limits of productivity depends on how efficiently a canopy converts fluctuating solar energy into biomass. Unlike controlled environments, irradiance in agricultural fields changes rapidly due to cloud cover, canopy architecture, and leaf movement (Long, Humphries and Falkowski, 1994; Ort, 2001). Under these dynamic conditions, plants frequently absorb more light energy than their metabolic capacity can accommodate, relying on the activation of photoprotective mechanisms to prevent irreversible damage to the photosynthetic apparatus (Demmig-Adams and Iii, 1992; Niyogi, 1999).

Plants dissipate excess energy primarily through non-photochemical quenching (NPQ) as heat. The effectiveness of this protection under fluctuating light depends on how closely NPQ dynamics synchronize with rapid changes in irradiance (Bassi and Dall’Osto, 2021; Kromdijk and Walter, 2023). Distinct NPQ components operate over different timescales, including rapidly relaxing energy-dependent quenching (qE), intermediate-relaxing components associated with zeaxanthin-dependent quenching and chloroplast movement (qM), and long-term quenching components commonly associated with photoinhibition (qI) (Li *et al*., 2002; Dall’Osto *et al*., 2014; Lam *et al*., 2026). Sustained high-light exposure promotes the accumulation of zeaxanthin through the violaxanthin–antheraxanthin–zeaxanthin (VAZ) xanthophyll cycle, whereas the slower reconversion of zeaxanthin under low light can delay NPQ relaxation (Demmig-Adams and Iii, 1992). Consequently, during rapid sun–shade transitions, excitation energy may continue to dissipate as heat instead of supporting carbon assimilation, reducing photosynthetic efficiency under fluctuating irradiance (Long *et al*., 2022).

Modeling analyses suggest that slow photoprotective recovery, alongside other dynamic constraints on photosynthesis, may reduce potential canopy CO₂ assimilation by 10-40% under fluctuating light (Long *et al*., 2022). In soybean (*Glycine max* Merr.), canopy simulations estimate daily reductions of approximately 13% in carbon assimilation due to slow NPQ relaxation (Wang *et al*., 2020), while natural variation among genotypes can cause differences of up to 1.6% in total carbon gain (Gotarkar *et al*., 2025). Targeted manipulation of these kinetic constraints has been demonstrated through the overexpression of Violaxanthin de-epoxidase (<u>V</u>DE), Photosystem II subunit S (<u>P</u>sbS) and <u>Z</u>eaxanthin epoxidase (ZEP) collectively referred to as VPZ. This strategy accelerated NPQ induction and relaxation and increased seed yield by 15% in tobacco and up to 33% in soybean under field conditions (Kromdijk *et al*., 2016; De Souza *et al*., 2022). These studies provide strong proof-of-concept for photosynthetic optimization. However, the translation of faster NPQ relaxation into higher yield appears to depend on finely tuned internal regulatory checkpoints that are not yet fully resolved (Singh *et al*., 2026). For instance, increasing PsbS levels can improve light-use efficiency and has also been linked to greater water-use efficiency under field conditions (Gotarkar *et al*., 2025). Conversely, uncoordinated acceleration of photoprotective relaxation can, in some cases, reduce biomass accumulation (Garcia-Molina and Leister, 2020). These observations suggest that the effectiveness of these photoprotective adjustments may depend on the stoichiometric balance between PsbS and xanthophyll cycle enzymes, as well as on the coordination between gene expression and pigment pool sizes across species and developmental stages.

Although bioengineering has revealed the potential of faster NPQ induction and relaxation to improve photosynthesis and crop yield (Kromdijk *et al*., 2016; De Souza *et al*., 2022), natural variation in photoprotective responses remains largely underexplored. Recent studies have begun to characterize this variation. In sorghum (*Sorghum bicolor*), large-scale analyses of field-grown diversity panels revealed substantial heritable variation in photoprotective traits and a complex genetic architecture underlying NPQ parameters (Vath *et al*., 2026). In soybean, analyses of the Nested Association Mapping (NAM) founder lines likewise identified significant diversity in NPQ relaxation (Gotarkar *et al*., 2025). However, while these studies reveal the genetic diversity underlying photoprotective responses, they do not address how modern crop breeding may have shaped these traits. In particular, it remains unclear whether a century of intensive selection for yield in soybeans has altered photoprotective dynamics such as NPQ relaxation.

We hypothesized that modern soybean genotypes would exhibit faster NPQ relaxation, accompanied by differences in VDE, PsbS, and ZEP transcript abundance, altered VAZ pigment dynamics, and improved photosynthetic performance under field conditions. To test this hypothesis, a historical panel of 24 soybean genotypes released between 1923 and 2021 was evaluated under field conditions. We characterized the relaxation dynamics of NPQ components and overall photosynthetic efficiency together with the molecular and biochemical factors underlying these processes. Specifically, we quantified transcript levels of *VDE*, *PsbS*, and *ZEP*, and profiled the VAZ xanthophyll cycle pigments (violaxanthin, antheraxanthin, and zeaxanthin). We conducted these measurements across distinct developmental stages to capture the transition from vegetative growth to reproductive stage. To assess how these photoprotective strategies influence productivity, we further correlated these physiological traits with seed number and weight.

## Materials and methods

### Plant material and experimental details

The study was conducted on a research farm at the University of Illinois Urbana-Champaign (40°02’ N, 88°14’ W, 228 m above sea level). Ten days before planting, the field was prepared by rototilling, cultivating, and harrowing. Seeds from 24 soybean genotypes released in different years were sown on May 27, 2025 (Table 1). Publicly developed genotypes were obtained from the USDA Soybean Germplasm Collection (Urbana, IL). The experiment followed a randomized complete block design with three blocks. Each block contained all genotypes randomly assigned to individual plots, totaling 72 plots. Each plot consisted of a single row oriented north-south, with 16 plants spaced 3.8 cm apart within the row, resulting in a row length of approximately 57.2 cm (Fig. S1). The experiment was rainfed, with limited supplemental manual irrigation around the V4 developmental stage to prevent severe drought stress.

**Table 1.** Soybean genotypes included in this study, with their year designation, maturity group, and Plant Introduction (PI) number. Year corresponds to cultivar release, germplasm registration, or approximate year of development/selection depending on genotype type.

| Genotype | Year | Maturity Group | PI Number |
| --- | --- | --- | --- |
| Dunfield | 1923 | III | PI 548318 |
| Illini | 1927 | II–III | PI 548348 |
| AK (Harrow) | 1928 | II | PI 548298 |
| Mandell | 1934 | III | PI 548381 |
| Adams | 1948 | III | PI 548502 |
| Ford | 1958 | III–IV | PI 548562 |
| Shelby | 1958 | III | PI 548574 |
| Ross | 1960 | III | PI 548612 |
| Adelphia | 1964 | II–III | PI 548503 |
| Wayne | 1964 | III | PI 548628 |
| Calland | 1968 | III | PI 548527 |
| Williams | 1971 | III | PI 548631 |
| Woodworth | 1974 | III | PI 548632 |
| Zane | 1984 | III–IV | PI 548634 |
| Thorne | 1993 | III | PI 564718 |
| Yale | 1994 | III | PI 584441 |
| Maverick | 1996 | III | PI 598124 |
| LD10-9168* | 2010 | III | - |
| LD11-2170* | 2011 | III | - |
| LD20-5413* | 2020 | III | - |
| LD20-11665* | 2020 | III | - |
| LD21-5547* | 2021 | III | - |
| BC4 Narrow Leaf** | 2021 | III | PI 612713A x LD11-2170 |
| BC4 Broad Leaf** | 2021 | III | PI 612713A x LD11-2170 |

\*LD lines are elite soybean breeding lines developed at the University of Illinois Urbana-Champaign (Cary and Diers, 2022).
\*\*BC4 Narrow Leaf and BC4 Broad Leaf are backcross-derived lines from LD11-2170 and PI 612713A with contrasting leaf morphology phenotypes (Tamang et al., 2023; 2025).
A dash indicates that no public PI number is available.

The average air temperature in Champaign, Illinois, during the experiment was 21.5°C. Maximum temperatures ranged from 26.2 °C to 34.7 °C, occurring in May and June 2025, respectively. Relative air humidity was lowest in June, while May had the lowest rainfall and July the highest. The average monthly accumulated solar radiation during the experiment was 536.2 MJ m⁻², with the highest value recorded in August (702.6 MJ m⁻²). Weather data were obtained from the SoyFACE Weather Station.

### NPQ relaxation measurements

The adjustment of NPQ during the transition from high to low light was evaluated using a modulated chlorophyll fluorescence imaging system (FluorCam FC800, Photon Systems Instruments, Drásov, Czech Republic). Samples were collected as subsamples within blocks (n = 3 blocks, 2 subsamples per block, 6 total observations per genotype). Six leaf disks (5.4 mm diameter) were collected from the uppermost fully expanded, sun-exposed leaf of each plant using a cork borer and treated as a single biological replicate. Samples were collected at the V5/V6 and R5/R6 developmental stages between 08:00 and 10:00 AM, following (Gotarkar *et al*., 2022). Leaf disks were placed facing down into 96-well plates and humidity was maintained by placing a half-wet nasal aspirator filter in each well (iHank-Nose B07P6XCTGV; Amazon, USA). Plates were sealed and wrapped in aluminum foil for overnight dark adaptation.

To induce NPQ, dark-adapted samples were illuminated for 10 min at 50 μmol m^2^ sec^-1^, followed by 15 min at 1600 μmol m^2^ sec^-1^, and a recovery phase of 45 min at 50 μmol m^2^ sec^-1^. NPQ kinetic phases were classified according to their relaxation behavior, including maximum NPQ (Max NPQ), defined as the peak NPQ reached during high-light illumination, fast-relaxing NPQ (qE; < 2 min), intermediate-relaxing NPQ (qM; 2–30 min), and slow-relaxing NPQ (qI; > 30 min). Parameters describing these phases included the amplitudes of each NPQ component (AqE and AqI), representing the magnitude of each relaxation phase, and the relaxation time constants (TqE and TqM), which describe the kinetics of NPQ relaxation. Background fluorescence was removed manually, and NPQ values were calculated using custom R and MATLAB scripts following the workflow described in (Gotarkar *et al*., 2022). Relaxation parameters were obtained by fitting a double-exponential decay function to NPQ values after the actinic light entered the recovery phase using the fit function in MATLAB R2018b, following (Dall’Osto *et al*., 2014)

### Leaf gas exchange measurements and LMA

The maximum CO₂ assimilation rate (*A*_max_) was determined under light- and CO₂-saturating conditions on the youngest fully expanded sun-exposed leaves using an infrared gas exchange system (LI-6800, LI-COR Environmental). Samples were collected as subsamples within blocks (n = 3 blocks, 2 subsamples per block, 6 total observations per genotype). Measurements were performed at two developmental stages (V5/V6 and R5/R6). To maintain high water potential and photosystem II efficiency, leaves were sampled predawn and petioles were recut underwater before measurements. Leaves were acclimated to1800 µmol m^−2^ s^−1^ photosynthetically active radiation (PAR) and 28 °C before the response of *A* to intercellular CO_2_ concentration (*Ci*) was measured. *A-Ci* curves were generated by measuring *A* at the following CO_2_ concentrations: 420, 300, 200, 100, 75, 50, 25, 420, 420, 600, 800, 1000, 1200, 1400, 1600, 1800 and 2000 mmol mol^-1^. *A*_max_ was calculated by fitting the gas exchange data to the models described by (Bellasio, Beerling and Griffiths, 2016). The maximum carboxylation rate of Rubisco (*V_cmax_*) and the maximum electron transport rate (*J*_max_) at 25 °C were estimated using the PhotoGEA R package (Lochocki, Salesse-Smith and McGrath, 2025). Leaf mass per area (LMA) was determined at R5/R6 using three leaf disks per plant (18 mm diameter) collected from fully expanded, sun-exposed leaves. Disk area was calculated based on known diameter, and samples were dried at 65 °C for 10 days prior to weighing. Two plants per block were sampled (n = 3 blocks, 2 subsamples per block, 6 total observations per genotype).

### Gene expression analysis

To link physiological variation with transcriptional responses, we classified genotypes into two groups based on NPQ relaxation. Genotypes such as Maverick, LD21-5547, Ross, and Shelby had low Max NPQ and slower relaxation (high TqE and TqM), whereas Mandell, Dunfield, BC4 Narrow Leaf, and Thorne had higher Max NPQ and faster relaxation (low TqE and TqM). These contrasting genotypes were selected for gene expression analyses targeting native gene paralogs involved in NPQ regulation and photoprotection, *VDE*, *PsbS*, and *ZEP*.

Leaf samples were collected at the R1/R2 growth stage between 11:30 AM and 12:30 PM. Three plants per block were collected (n = 3 blocks, 3 subsamples per block, 9 total observations per genotype). From each plant, three leaf disks (13.4 mm diameter) were harvested from the youngest fully expanded sun-exposed leaf using a cork borer (H-9663; Humboldt, USA). Samples were immediately placed in 2 mL nuclease-free tubes, flash frozen in liquid nitrogen, and stored at –80 °C. For homogenization, samples were ground using one 4 mm grinding bead (2150, Cole-Parmer, USA) at 20 Hz for 1.5 min (Tissue Lyser, Qiagen, Germany), repeated three times, with samples kept in liquid nitrogen between cycles. RNA was extracted using the RNeasy Plant Kit (74904, Qiagen, Germany) with minor modifications to the manufacturer’s instructions. After addition of Buffer RLT, a guanidine thiocyanate-containing lysis buffer used to inactivate RNases, samples were centrifuged at 12,000 × g for 1 min. An on-column DNase treatment was then performed following the RW1 wash buffer step. Briefly, 350 µL of RW1 was added and centrifuged, followed by the application of 80 µL DNase I (79254, Qiagen, Germany) directly onto the spin column membrane and incubation for 15 min at room temperature. An additional 350 µL of RW1 was subsequently applied, and the number of washes with RPE wash buffer (Qiagen) was increased from two to four. RNA quantity and quality were assessed using a NanoDrop One spectrophotometer (Thermo Fisher, USA). cDNA was synthesized using the SuperScript™ III First-Strand Synthesis System (18080051, Thermo Fisher, USA) with random hexamers. DNA contamination was evaluated and identified using a set of no reverse transcript (NRT) control samples. An additional DNase treatment was applied to the RNA using ezDNase™ (11766051, Thermo Fisher, USA) at half the recommended reaction volume, with the RNA input reduced proportionally to maintain the same DNase-to-RNA ratio as specified by the manufacturer, allowing the reaction volume to be used directly in a new cDNA synthesis as described above. Evaluation of the cDNA NRT samples from the twice DNase treated RNA showed no DNA contamination. The resulting cDNA was diluted 1:100 in Milli-Q water. Two *VDE* gene paralogs (VDE1 - *Glyma.03G253500,* VDE2 - *Glyma.19G251000*), three *ZEP* paralogs (*ZEP1* - *Glyma.17G174500*, *ZEP2* -*Glyma.11G055700*, *ZEP3* - *Glyma.09G000600*), and two *PsbS* paralogs (*PSBS1* - *Glyma.04G249700, PSBS2 - Glyma.06G113200*) were analyzed.

The reference genes were *GmELF1* (*Glyma.02G276600*) and *GmCYP* (*Glyma.12G024700*) (De Souza *et al*., 2022). qPCR reactions were performed using either iTaq Universal SYBR® Green Supermix (1725120, Bio-Rad, USA) or SYBR® Green Supermix (1725270, Bio-Rad, USA) with 4 µL cDNA in 384-well plates on a QuantStudio 7 Pro system (Applied Biosystems, USA). The cycling program consisted of 95 °C for 2 min, followed by 40 cycles of 95 °C for 15 s and a gene-specific annealing temperature for 30 s. Primer sequences and annealing temperatures are provided in Table S1.

Primers were designed based on *Glycine max* reference sequences (Phytozome v12.1) following MIQE guidelines. Specificity was assessed using NCBI Primer-BLAST, and amplicon sizes ranged from 70 to 150 bp. Amplification efficiency and linear dynamic range were determined using standard curves, and only primers with efficiencies between 90% and 110% were used. Additional primer characteristics are provided in Tables S1 and S2. Calibrated normalized relative quantities (CNRQ) were calculated using qbase+ version 3.2 (CellCarta, Canada) following (Hellemans *et al*., 2007).

### Xanthophyll pigment analysis

To assess whether differences in NPQ relaxation were reflected at the pigment level, we analyzed the same contrasting genotype groups used for gene expression analysis. Samples were collected between 11:30 AM and 12:30 PM at the R1/R2 developmental stage. Three plants per block were sampled (n = 3 blocks, 3 subsamples per block, 9 total observations per genotype). From each plant, three leaf disks (13.4 mm diameter) were excised from the youngest fully expanded sun-exposed leaf using a cork borer (H9663; Humboldt, USA). Samples were flash-frozen in liquid nitrogen immediately after collection and stored at –80 °C until processing. The frozen material was ground in 2 mL tubes containing two 4 mm stainless steel beads using a tissue lyser at 24 Hz for 1.5–2 min, with the procedure repeated once after inverting the adapters. Pigments were extracted from the resulting pellets using two successive additions of 100 μL of 100% acetone (Leonelli, 2022). Pigment composition was subsequently separated and quantified according to (Müller-Moulé, Conklin and Niyogi, 2002) using an Agilent 1290 Infinity UPLC system (Agilent, Santa Clara, CA) equipped with a Spherisorb 5 μm ODS1 column (Waters Corp., Milford, MA, USA). A quaternary pump provided a linear gradient starting at 14% solvent A (0.1 M Tris-HCl pH 8.0), 84% solvent B (acetonitrile), and 2% solvent C (methanol) for 15 min; transitioning to 68% solvent C and 32% solvent D (ethyl acetate) for 4 min; and finally returning to the initial solvent composition for 6 min at a flow rate of 1.2 mL min⁻¹. Chromatograms were manually integrated using Control Panel for Agilent OpenLab software. Pigment concentrations were normalized to total chlorophyll content.

### Seed traits

All plants within each plot were harvested manually. Pods were threshed, and seeds from each plant were collected separately. Seed weight per plant (g) and seed number per plant were measured at the individual plant level.

### Statistical analysis

Normality and homogeneity of variances were assessed using the Shapiro–Wilk and Levene’s tests, respectively. When assumptions of normality and homoscedasticity were met, differences among genotypes were analyzed using one-way analysis of variance (ANOVA), followed by Tukey’s post hoc HSD test. For group comparisons, genotypes were classified into two groups: the seven oldest (1923–1958) and the seven most recently released genotypes (2010–2021) (Table 1). Group means were compared using Welch’s two-sample t-test.

Relationships between variables were evaluated using Spearman’s rank correlation based on genotype-level means. Correlation coefficients (ρ) were calculated, and P-values were adjusted for multiple comparisons using the Benjamini–Hochberg false discovery rate (FDR), applied within each developmental stage. Linear regression lines shown in figures are for visualization purposes. When necessary, data were transformed prior to analysis to meet model assumptions. Statistical significance was defined as P < 0.05. Statistical tests applied to each dataset are reported in the corresponding figure or table legends. All analyses were conducted in R (R Core Team, 2024) and SigmaPlot (Version 13, 2026, Systat Software, Inc., San Jose, California).

## Results

### NPQ relaxation has no consistent relationship with year of release

To assess whether NPQ relaxation traits have changed with breeding, we calculated Spearman’s rank correlations between NPQ parameters and year of release (YOR) (Fig. 1). At the V5/V6 stage, none of the five parameters (Max NPQ, TqE, TqM, AqE, AqI) were significantly associated with YOR (all P > 0.05; Fig. 1A). At R5/R6, AqE was the only exception, showing a negative correlation with YOR (ρ = −0.60, P = 0.01; Fig. 1B).

**Fig. 1.**
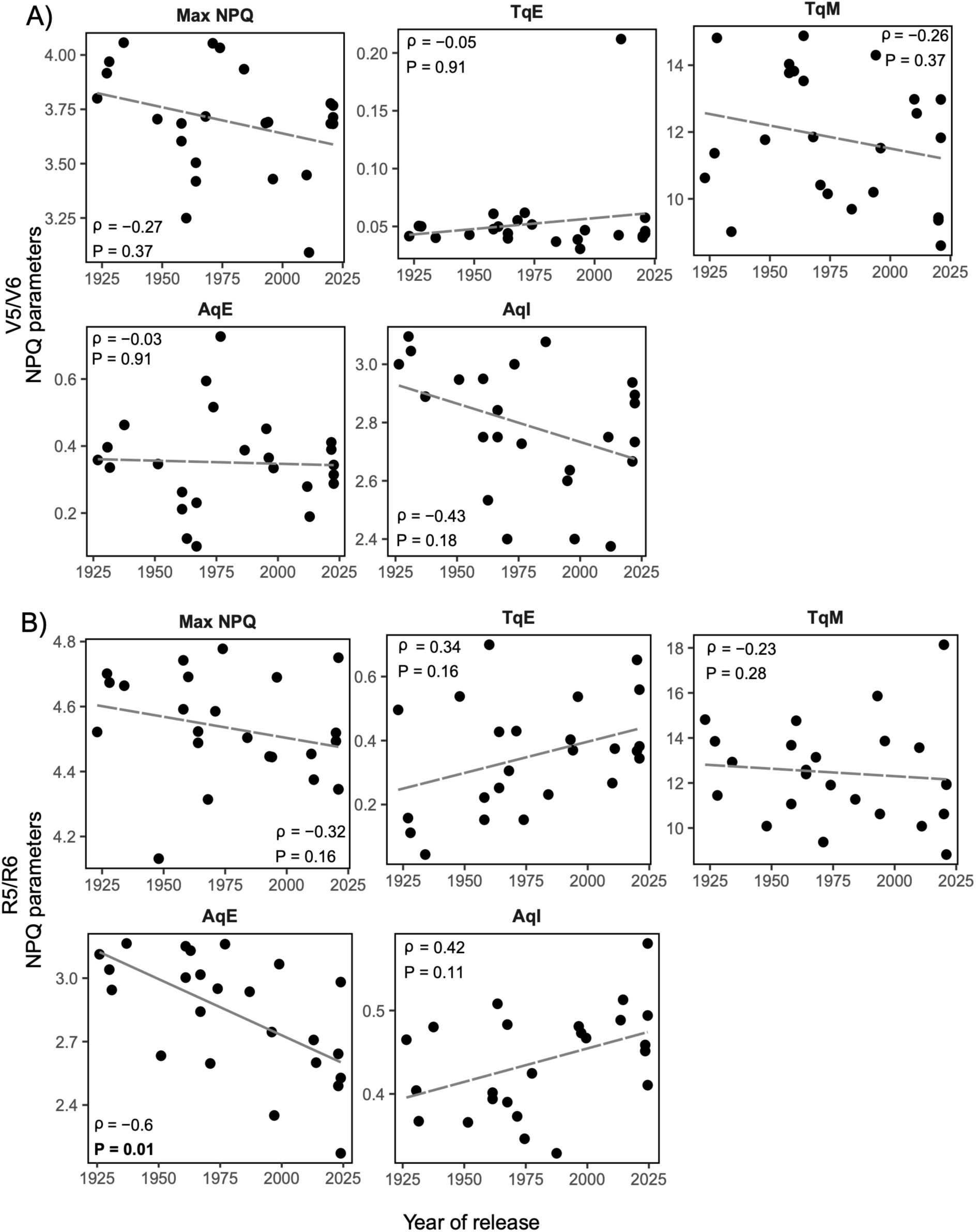
Correlations between non-photochemical quenching (NPQ) relaxation parameters and year of release (YOR) in different soybean cultivars. Scatter plots show correlations between each NPQ parameter (Max NPQ, TqE, TqM, AqE, and AqI) and YOR at the V5/V6 (A) and R5/R6 (B) developmental stages. Values represent the mean (± SE) of three replicate blocks (*n* = 3) with two subsamples. Lines show linear fits for visualization only; correlations were assessed using Spearman’s rank correlation. Spearman correlation coefficients (ρ) and FDR-adjusted *P*-values (Benjamini–Hochberg correction, applied within each developmental stage) are shown in each panel. Dashed lines indicate non-significant correlations. **Alt text:** Scatter plots showing relationships between soybean genotype release year and NPQ relaxation parameters at vegetative (top two rows) and reproductive (bottom two rows) developmental stages. Panels include maximum NPQ (Max NPQ), relaxation time constants (TqE and TqM), and amplitudes of NPQ components (AqE and AqI). Most parameters showed no significant association with genotype release year, except AqE during the reproductive stage, which showed a significant negative correlation. Solid lines indicate significant relationships and dashed lines indicate non-significant relationships.

### NPQ kinetic parameters varies across genotypes and developmental stages

NPQ relaxation varied across genotypes at both the V5/V6 and R5/R6 stages, with no clear pattern across the measured NPQ parameters. This lack of structure was also evident when comparing the seven oldest genotypes (Dunfield, Illini, AK (Harrow), Mandell, Adams, Ford, and Shelby), released between 1923 and 1958, with the seven most recently released genotypes (LD10-9168, LD11-2170, LD20-11665, LD20-5413, BC4 Broad Leaf, BC4 Narrow Leaf, and LD21-5547), released between 2010 and 2021 representing the extremes of the release range (Fig. 2 and Table 1). Differences were also observed within genotypes across developmental stages. Some genotypes maintained similar Max NPQ values across stages, whereas others showed greater variation in relaxation-related parameters, particularly TqE and TqM. For example, LD10-9168, LD11-2170, and LD20-5413 had relatively low Max NPQ at both stages but differed in other parameters. In LD11-2170, TqE was high at V5/V6 but strongly reduced at R5/R6 (Fig. 2). Consistent with these observations, Max NPQ showed less variation among genotypes than TqE and TqM at both developmental stages (Table 2).

**Fig. 2.**
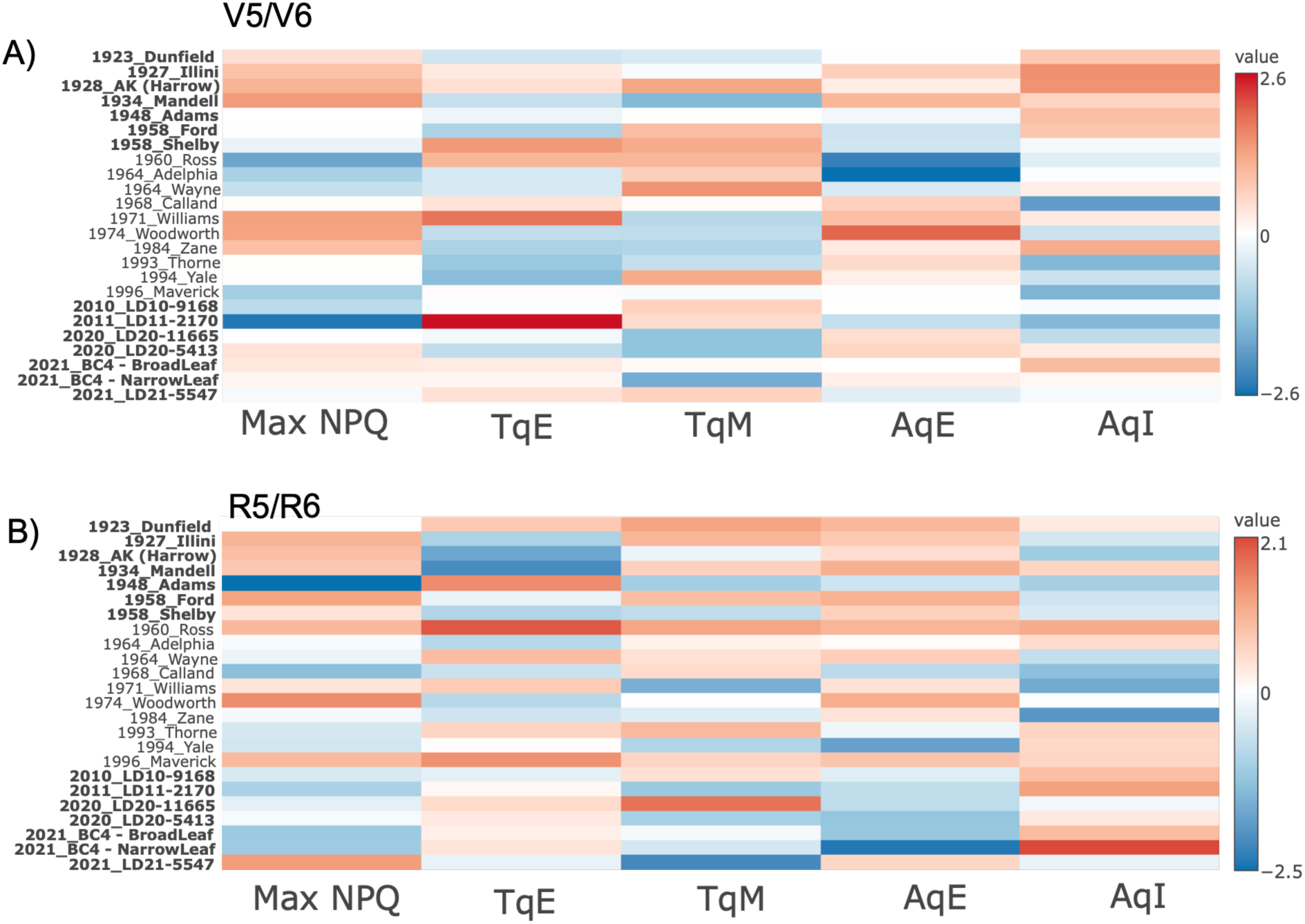
Variation in non-photochemical quenching (NPQ) relaxation dynamics in soybean. Heatmaps show variation in AqI, Max NPQ, TqE, and TqM at the V5/V6 (A) and R5/R6 (B) developmental stages. Data are means (± SE) of three replicate blocks (*n* = 3), each with two subsamples. Values were log₁₀-transformed and autoscaled (Z-scores). Rows represent genotypes and columns represent NPQ parameters. The color scale indicates relative values (red = high, blue = low). Genotypes shown in bold represent the seven oldest (1923–1958) and the seven most recently released genotypes (2010–2021). **Alt text:** Heatmaps showing normalized NPQ relaxation parameters across a historical panel of soybean genotypes at vegetative (top panel) and reproductive (bottom panel) developmental stages. Rows represent individual genotypes ordered by year of release, and columns represent NPQ parameters including Max NPQ, TqE, TqM, AqE, and AqI. Color intensity indicates relative parameter values, ranging from lower values in blue to higher values in red. No clear directional trend across genotype release years is apparent for most NPQ parameters at either developmental stage.

**Table 2.** Summary of NPQ relaxation dynamics for 24 soybean genotypes based on mean values across all samples.

| Parameter | V5/V6 |  |  | R5/R6 |  |  |
| --- | --- | --- | --- | --- | --- | --- |
|  | Range | Median | Variation (%) | Range | Median | Variation (%) |
| Max NPQ | 3.09 - 4.0 | 3.70 | 31.23 | 4.12 - 4.77 | 4.52 | 15.63 |
| TqE | 0.03 - 0.06 | 0.05 | 102.03 | 0.04 - 0.69 | 0.36 | 1502.11 |
| TqM | 8.59 - 14.88 | 11.80 | 73.04 | 8.82 - 18.13 | 12.17 | 105.65 |
| AqE | 0.1 - 0.7 | 0.34 | 621.24 | 2.1 - 3.1 | 2.9 | 45.6 |
| AqI | 2.45 - 3.0 | 2.78 | 23.88 | 0.32 - 0.5 | 0.45 | 76.5 |

### Modern genotypes show higher TqE and lower TqM, AqE, and Max NPQ

For this and subsequent analyses, genotypes were classified as old or modern based on release year, representing the seven oldest and seven most recently released genotypes (Fig. 3). Across both developmental stages, modern genotypes showed higher TqE and lower TqM, AqE, and Max NPQ than old genotypes. In contrast, AqI was lower at V5/V6 but higher at R5/R6.

**Fig. 3.**
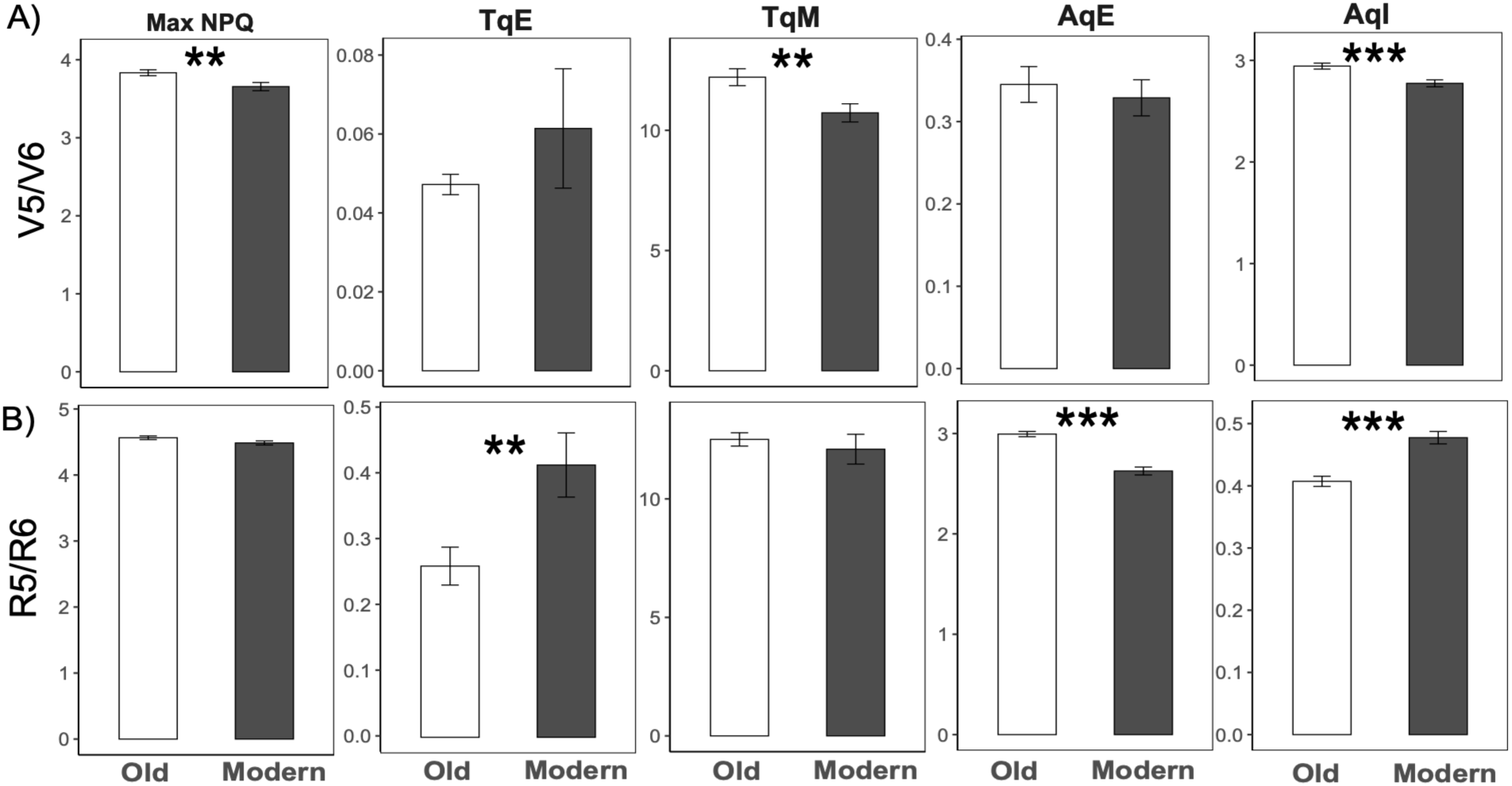
Comparison of non-photochemical quenching (NPQ) relaxation parameters between old and modern soybean genotypes. Data are means ± SE of three replicate blocks (*n* = 3), each with two subsamples. Bar plots show Max NPQ, TqE, TqM, AqE, and AqI for the seven oldest and seven most recently released genotypes at the V5/V6 (A) and R5/R6 (B) developmental stages. The genotypes included in each group are listed in Table 2. Asterisks indicate significant differences between groups based on Welch’s two-sample t-test (*P < 0.05, **P < 0.01, ***P < 0.001). **Alt text:** Bar plots comparing NPQ relaxation parameters between old and modern soybean genotypes at vegetative (top row) and reproductive (bottom row) developmental stages. Panels include maximum NPQ (Max NPQ), relaxation time constants (TqE and TqM), and amplitudes of NPQ components (AqE and AqI). Bars represent mean values with error bars indicating standard error. Significant differences between old and modern genotypes are indicated by asterisks. Modern genotypes generally showed lower Max NPQ, TqM, and AqE values, and higher AqI values, particularly during the reproductive stage.

### Photosynthetic capacity is higher in modern genotypes

To complement the analysis of NPQ relaxation parameters, we quantified key biochemical indicators of photosynthetic capacity (*A*_max_, *V_cmax_* and *J*_max_) at V5/V6 and R5/R6 stages. Across both stages, all three parameters were greater in modern genotypes than in older genotypes (Fig. 4). Genotype-level variation is shown in detail in Fig. S2. In contrast, leaf mass per area (LMA) did not differ among genotypes (Fig. S3).

**Fig. 4.**
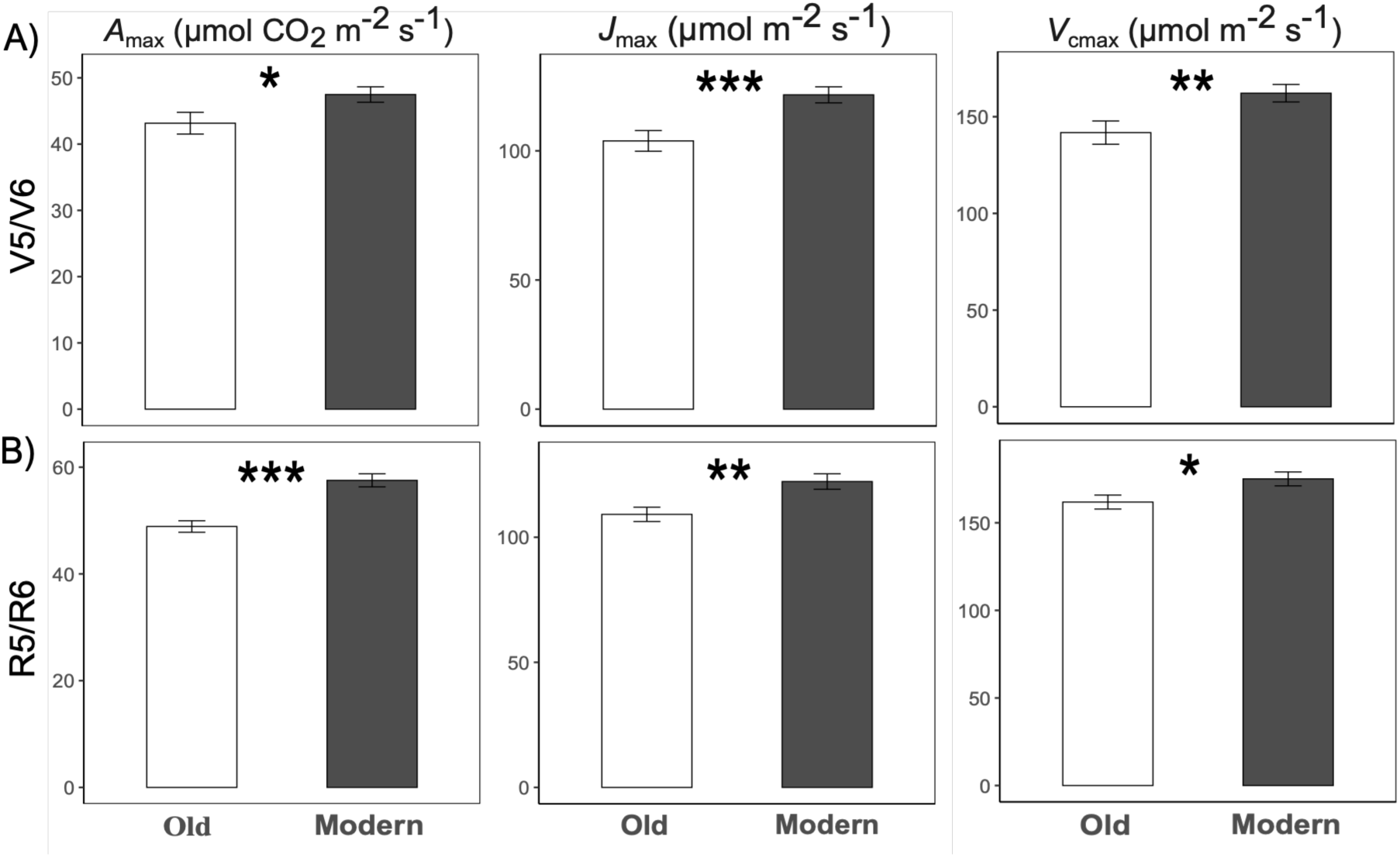
Differences in photosynthetic capacity parameters between old and modern soybean genotypes. Data are means ± SE of three replicate blocks (*n* = 3), each with two subsamples. Bar plots show *A*_max_, *V_cmax_* and *J*_max_ for the seven oldest and seven most recently released genotypes at the V5/V6 (A) and R5/R6 (B) developmental stages. Genotypes included in each group are listed in Table 2. Asterisks indicate significant differences between groups based on Welch’s two-sample t-test (*P < 0.05, **P < 0.01, ***P < 0.001). **Alt text:** Bar plots comparing photosynthetic parameters between old and modern soybean genotypes at vegetative (top row) and reproductive (bottom row) developmental stages. Panels show maximum net CO₂ assimilation rate (*A*_max_), maximum electron transport rate (Jmax), and maximum carboxylation rate of Rubisco (*V*_cmax_). Bars represent mean values with error bars indicating standard error. Significant differences between old and modern genotypes are indicated by asterisks. Modern genotypes generally exhibited higher photosynthetic capacity than old genotypes, particularly during the reproductive stage.

### Photosynthetic capacity increases with year of release

We evaluated correlations between photosynthetic capacity parameters and YOR using Spearman’s rank correlation (Fig. 5). At the V5/V6 stage, *A*_max_, *V_cmax_ and J*_max_ were positively but non-significantly correlated with YOR. At the R5/R6 stage, all three parameters were positively and significantly correlated with YOR: *A*_max_ (ρ = 0.65, P = 0.002), *J*_max_ (ρ = 0.46, P = 0.022), and *V_cmax_* (ρ = 0.50, P = 0.019).

**Fig. 5.**
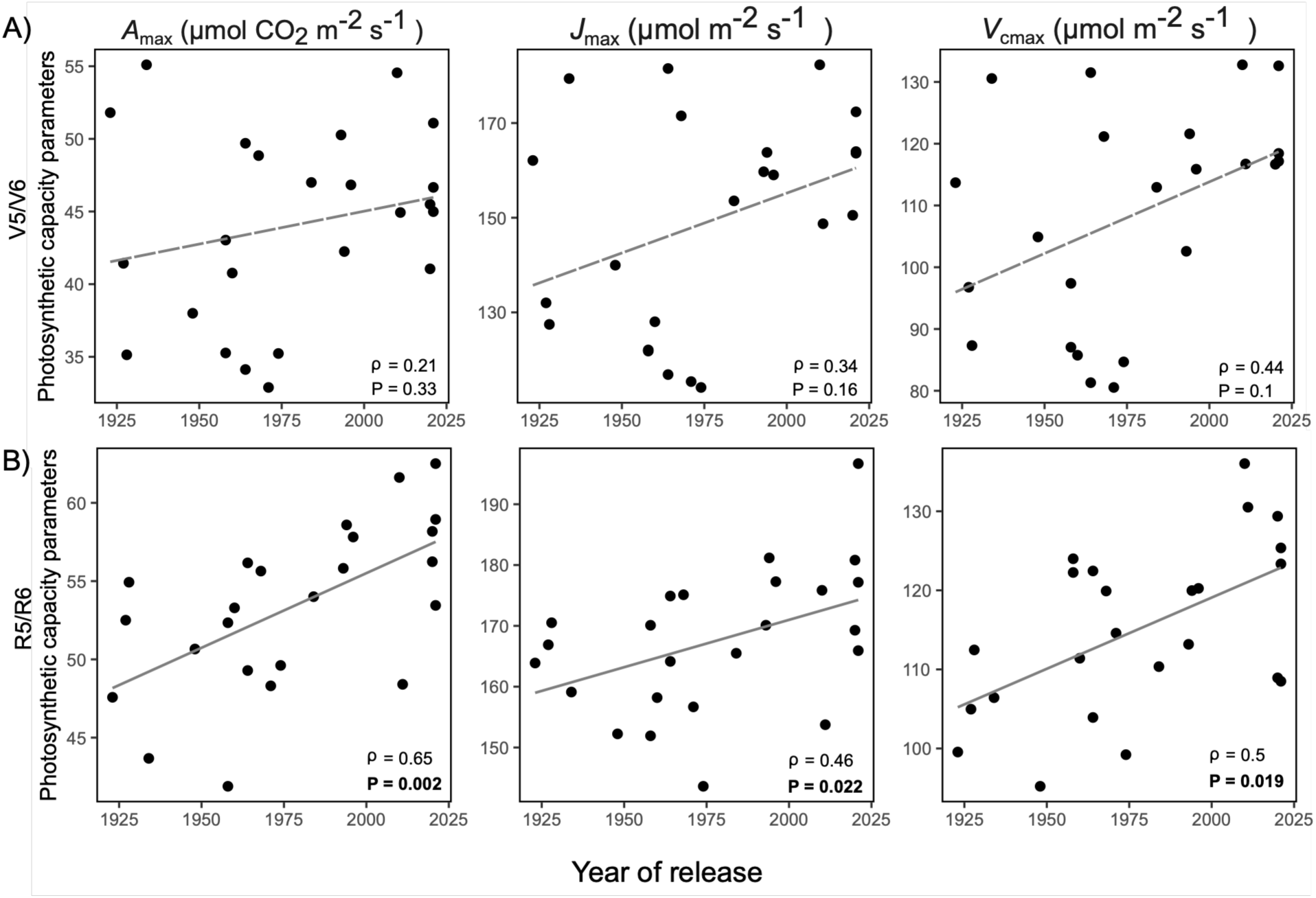
Correlations between photosynthetic capacity parameters and year of release (YOR) in soybean genotypes. Scatter plots show correlations between each parameter (*A*_max_, *J*_max_, and *V*_cmax_) and YOR at the V5/V6 (A) and R5/R6 (B) developmental stages. Values represent the mean (± SE) of three replicate blocks (*n* = 3) with two subsamples. Lines show linear fits for visualization only; correlations were assessed using Spearman’s rank correlation. Spearman correlation coefficients (ρ) and FDR-adjusted *P*-values (Benjamini–Hochberg correction, applied within each developmental stage) are shown in each panel. Dashed lines indicate non-significant correlations. **Alt text:** Scatter plots showing relationships between soybean genotype release year and photosynthetic capacity parameters at vegetative (top row) and reproductive (bottom row) developmental stages. Panels include maximum net CO₂ assimilation rate (*A*_max_), maximum electron transport rate (*J*_max_), and maximum carboxylation rate of Rubisco (*V*_cmax_). Most photosynthetic parameters showed positive associations with genotype release year, particularly during the reproductive stage, where modern genotypes exhibited greater photosynthetic capacity. Solid lines indicate significant relationships and dashed lines indicate non-significant relationships.

### Modern genotypes show greater seed production

The mean number of seeds per plant and mean seed weight per plant were higher in modern genotypes than in older genotypes (Fig. 6). Seed weight ranged from 6 to 17 g per plant in old genotypes and from 12 to 22 g per plant in modern genotypes. The number of seeds per plant ranged from 48 to 122 in old and from 100 to 199 seeds per plant in modern genotypes.

**Fig. 6.**
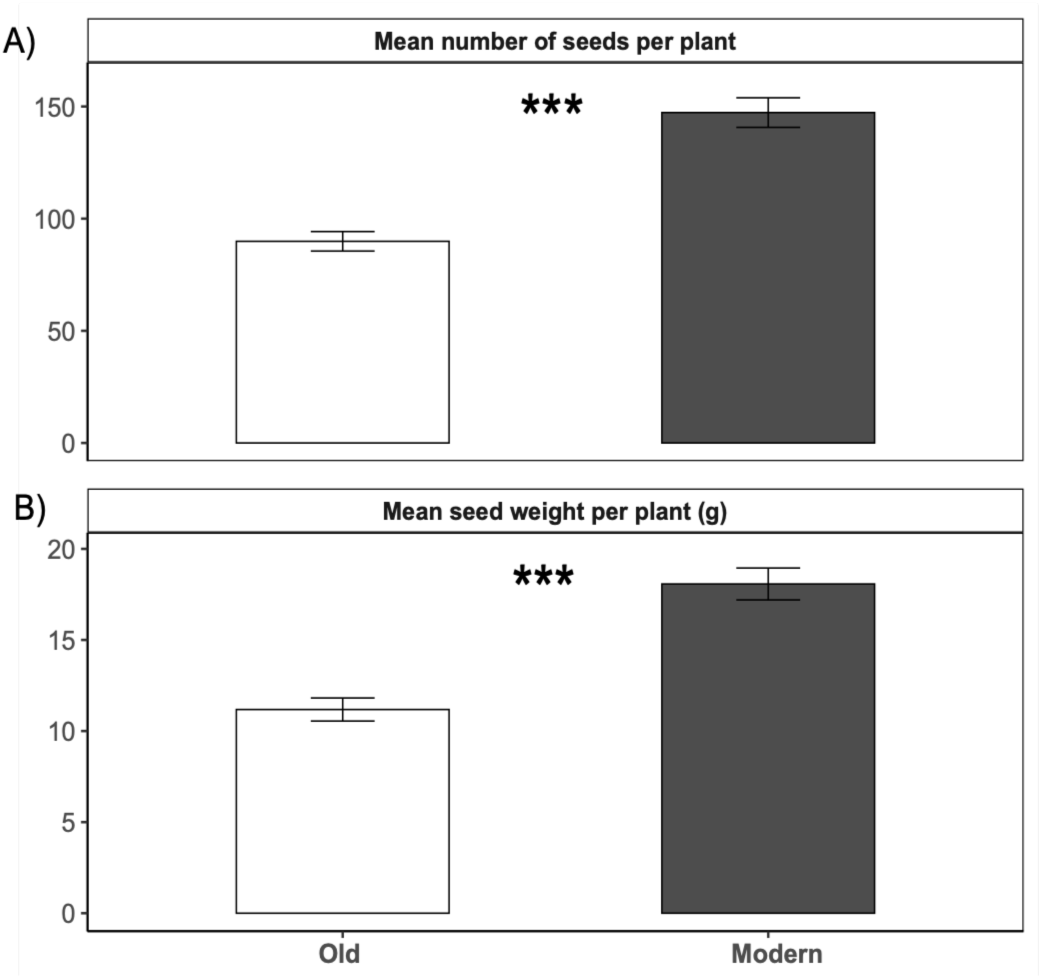
Seed number and seed weight in old and modern soybean genotypes. (A) Mean number of seeds per plant. (B) Mean seed weight per plant (g). Data are means ± SE for the seven oldest and seven most recently released genotypes. Genotypes included in each group are listed in Table 2. Asterisks indicate significant differences between groups based on Welch’s two-sample t-test (*P < 0.05, **P < 0.01, ***P < 0.001). **Alt text:** Bar plots comparing mean seed number per plant (top panel) and mean seed weight per plant (bottom panel) between older and modern soybean genotypes. Bars represent mean values with error bars indicating standard error. Modern genotypes exhibited significantly greater seed number and seed weight compared to older genotypes, as indicated by asterisks.

### Photosynthetic parameters (A_max_ and V_cmax_), but not NPQ relaxation traits, were positively correlated with seed weight and number

We next assessed Spearman correlations between seed traits (seed number and weight) and each NPQ relaxation or photosynthetic capacity parameter (Fig. 7). At V5/V6, most NPQ-related parameters were not significantly associated with seed traits, except AqI, which was negatively correlated with seed number (ρ = −0.42, P = 0.04; Fig. S4). At R5/R6, AqI was positively correlated with seed number, whereas AqE was negatively correlated with both seed number and weight. In contrast, photosynthetic parameters (*A*_max_ and *V_cmax_*) were consistently and positively correlated with both seed number and seed weight during the reproductive stage. (Fig. 7).

**Fig. 7.**
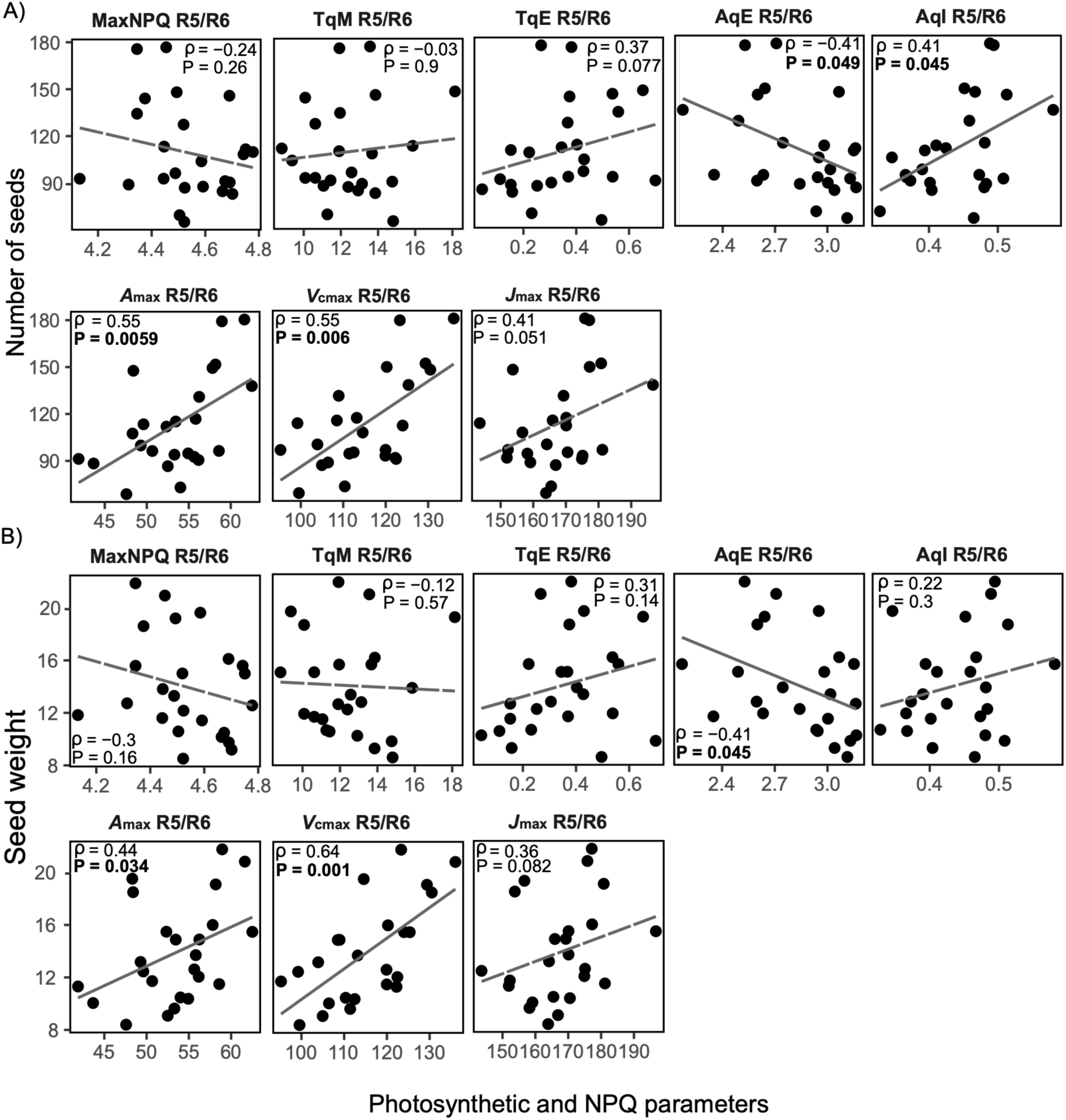
Scatter plots show relationships between NPQ parameters (Max NPQ, TqE, TqM, AqE, AqI), photosynthetic capacity (*A*_max_, *J*_max_ and *V*_cmax_), and seed traits, seed number (A) and seed weight (B) per plant at the R5/R6. Values represent the mean (± SE) of three replicate blocks (*n* = 3) with two subsamples. Lines show linear fits for visualization only; correlations were assessed using Spearman’s rank correlation. Spearman correlation coefficients (ρ) and FDR-adjusted *P*-values (Benjamini–Hochberg correction, applied within each developmental stage) are shown in each panel. Dashed lines indicate non-significant correlations. **Alt text:** Scatter plots showing relationships between seed traits and physiological parameters in soybean genotypes at the R5/R6 developmental stage. Panels compare seed number per plant (A) and seed weight per plant (B) with NPQ relaxation parameters, including Max NPQ, TqE, TqM, AqE, and AqI, as well as photosynthetic capacity parameters *A*_max_, *J*_max_, and *V*_cmax_. Each point represents the mean value for an individual genotype, with dashed lines showing linear fits for visualization. Most NPQ parameters showed weak or non-significant relationships with seed traits, whereas *A*_max_ and *V*_cmax_ were positively associated with seed number and seed weight. AqE showed negative correlations with both seed traits, while AqI was positively correlated with seed number. Spearman correlation coefficients and adjusted P-values are shown within each panel.

### Pigment composition and gene expression do not fully explain NPQ relaxation differences

To explore the molecular basis of NPQ relaxation, we selected eight genotypes with contrasting extremes of relaxation kinetics (Fig. S5): four slow-relaxing genotypes (Maverick, LD21-5547, Ross, and Shelby) and four fast-relaxing genotypes (Mandell, Dunfield, BC4 Narrow Leaf, and Thorne). VDE, PsbS, and ZEP belong to multigene families in soybean and likely contribute collectively to NPQ regulation. Expression values of all identified paralogs were therefore summed to estimate total family-level expression. Individual paralog expression profiles are presented in Fig. S6, whereas individual xanthophyll-cycle pigments (violaxanthin, antheraxanthin, and zeaxanthin) are shown in Fig. S7. Figure 8 summarizes the total xanthophyll-cycle pigment pool (VAZ), de-epoxidation state (VAZ.DES) reflecting the relative conversion of violaxanthin to zeaxanthin, total family-level expression of VDE, PsbS, and ZEP, and the VDE/ZEP expression ratio.

**Fig. 8.**
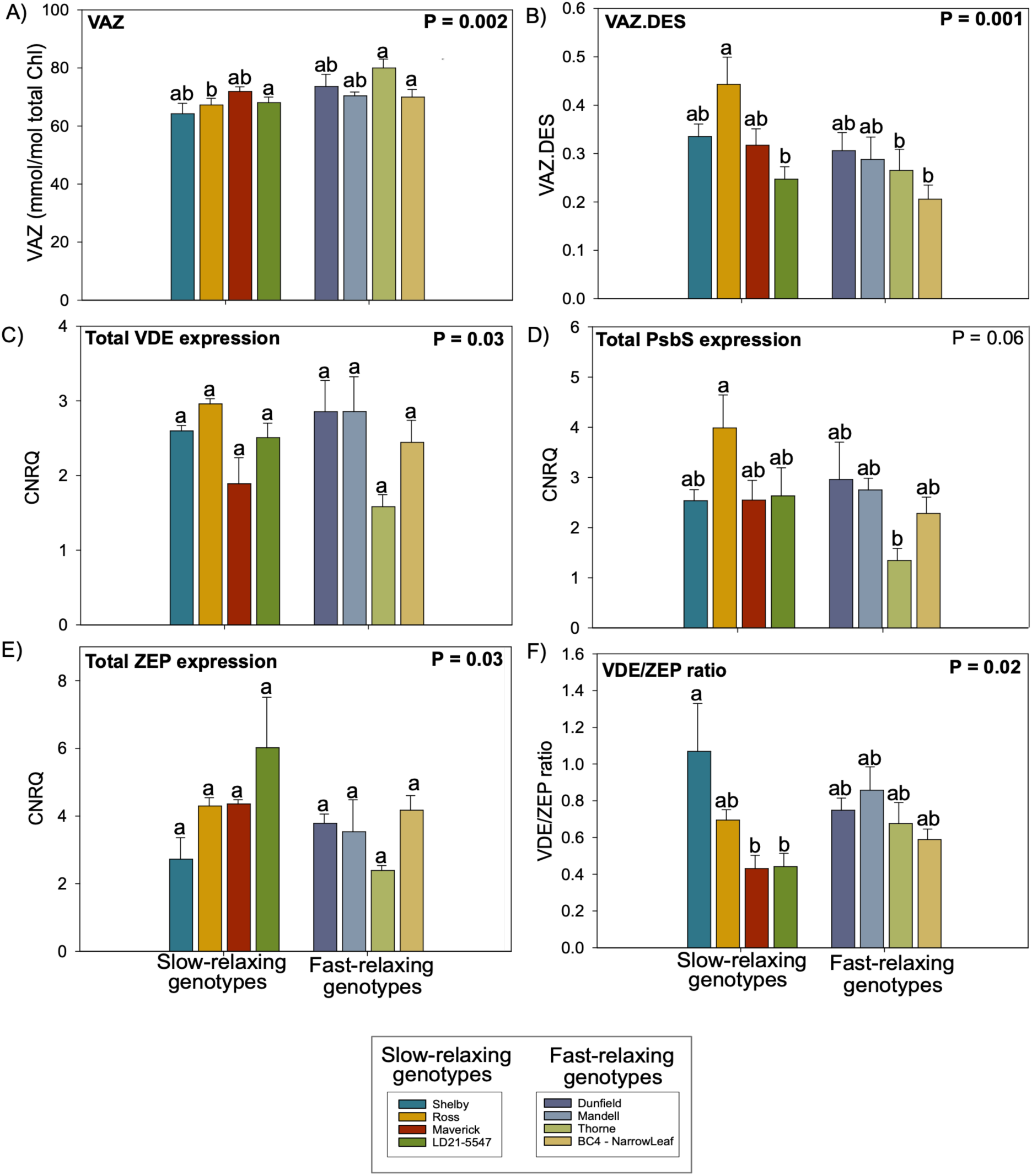
Pigment composition and expression of NPQ-related genes in selected soybean genotypes. Data are means ± SE of three replicate blocks (*n* = 3), each with three subsamples. Bar plots show the total xanthophyll-cycle pigment pool (VAZ; V + A + Z) (A), de-epoxidation state (VAZ.DES) (B), and total relative expression of VDE (C), PsbS (D), and ZEP (E), calculated as the sum of the all paralogs identified for each gene family, as well as the VDE/ZEP expression ratio (F). VAZ.DES was calculated as (Zea + 0.5 × Ant) / (Vio + Ant + Zea), with pigment pools normalized to total chlorophyll (mmol mol⁻¹). Gene expression is shown as calibrated normalized relative quantity (CNRQ) values. Letters indicate significant differences among genotypes (one-way ANOVA followed by Tukey’s post hoc test, P < 0.05). P-values are shown in each panel. **Alt text:** Bar plots comparing xanthophyll-cycle pigment composition and expression of NPQ-related genes among soybean genotypes classified as slow-relaxing or fast-relaxing NPQ types. Panels show total xanthophyll-cycle pigment pool (VAZ), xanthophyll de-epoxidation state (VAZ.DES), total VDE expression, total PsbS expression, total ZEP expression, and the VDE/ZEP expression ratio. Bars represent mean values with error bars indicating standard error. Significant differences among genotypes are indicated by letters above bars, and P-values are shown within panels. Variation in pigment composition and gene expression overlapped between fast- and slow-relaxing genotypes, indicating genotype-specific patterns rather than a consistent separation between relaxation groups.

Significant genotypic differences were detected for VAZ (P = 0.002), VAZ.DES (P = 0.001), total VDE expression (P = 0.03), total ZEP expression (P = 0.03), and the VDE/ZEP expression ratio (P = 0.02), whereas total PsbS expression did not differ significantly among genotypes (Fig. 8). Despite these differences, pigment composition and gene expression overlapped between fast- and slow-relaxing genotypes, indicating genotype-specific variation. For example, Ross, a slow-relaxing genotype, showed lower total VAZ content (Fig. 8A) but a higher de-epoxidation state of the xanthophyll cycle (Fig. 8B) and higher total PsbS expression (Fig. 8D) than Thorne, a fast-relaxing genotype.

## Discussion

Soybean yields in the United States increased from 1648 kg ha⁻¹ (24.5 bu ac⁻¹) in 1965 to more than 3500 kg ha⁻¹ (>50 bu ac⁻¹) in recent years (USDA-NASS, 2025). These gains have been attributed to improvements from breeding and agronomic practices (Anderson *et al*., 2019). However, despite this sustained increase in productivity, our results show no consistent directional trend in NPQ relaxation with YOR (Fig. 1). Instead, NPQ parameters varied widely among genotypes at both developmental stages (Fig. 2; Table 2), indicating considerable diversity in photoprotective responses without a clear pattern of improvement through breeding. Previous work in soybean reported similar results, showing that NPQ parameters do not follow consistent developmental trends and are strongly influenced by environmental conditions, with no clear relationship to yield across cultivar diversity (Gotarkar *et al*., 2025). Significant differences across all relaxation parameters (Table 2) further indicate that soybean retains a broad range of photoprotective responses that remain largely untapped for crop improvement.

Similar levels of variation have been reported in sorghum (Vath *et al*., 2026) and soybean nested association mapping populations (Gotarkar *et al*., 2025), highlighting both genetic and environmental contributions to NPQ relaxation. In sorghum, this variation has been linked to a complex polygenic architecture. In soybean, environmental factors such as temperature and vapor pressure deficit explain 8-47% of the variation in NPQ relaxation, depending on the parameter and growing season (Gotarkar *et al*., 2025). Plant development also contributes to this variation. As soybean shifts from vegetative biomass accumulation to reproductive seed filling, stronger sink demand for carbohydrates may alter the balance between photochemical energy use and dissipation (Park *et al*., 2023). Nitrogen remobilization from Rubisco and light-harvesting complexes to developing seeds may further affect this balance by changing the metabolic demand on the photosynthetic machinery (Paul and Foyer, 2001). Consistent with this interpretation, previous studies have shown that NPQ engagement and relaxation vary with leaf developmental stage, canopy position, and structural changes in the light environment (Jiang *et al*., 2005; Bielczynski *et al*., 2017; Foo *et al*., 2020).

Grouping genotypes into old and modern categories revealed consistent differences despite the lack of a directional trend across release years. Modern genotypes had higher TqE and lower TqM, AqE, and Max NPQ across developmental stages (Fig. 3) Higher TqE indicates slower relaxation of the fast NPQ component, whereas lower TqM suggests faster relaxation of intermediate components. Reduced AqE and Max NPQ also indicate lower overall engagement of photoprotective energy dissipation. Together, these results suggest that modern genotypes differ from older genotypes not through uniform improvement in NPQ performance, but through shifts in the relative contribution and relaxation behavior of NPQ components.

Correlation analyses showed no consistent overall relationship between NPQ parameters and seed number or seed weight (Fig. 7). Among the five parameters analyzed, only AqE and AqI were significantly associated with seed traits at R5/R6, and these relationships showed opposite patterns. AqE was negatively correlated with both seed number and seed weight, whereas AqI was positively correlated with seed number. These contrasting relationships indicate that NPQ should not be interpreted as a single functional trait when linking photoprotection to reproductive performance. Instead, different NPQ components reflect distinct physiological processes that operate on different timescales and are controlled by different regulatory mechanisms (Murchie and Niyogi, 2011). As a result, they may show independent or even opposing relationships with seed traits. Previous work in soybean also showed that NPQ components differ in their stability and responsiveness across environments, with some parameters remaining consistent across years and others showing greater plasticity (Gotarkar *et al*., 2025). In this context, the opposing associations observed here for AqE and AqI likely reflect component-specific responses rather than a unified effect of NPQ on seed traits. Consistent with this interpretation, modern genotypes showed lower AqE but higher AqI than older genotypes at the same developmental stage (Fig. 3). This pattern reinforces that NPQ components did not change in a coordinated manner with breeding. Instead, breeding appears to have shifted the balance between rapid and sustained photoprotective processes.

In contrast to NPQ traits, photosynthetic parameters, including *A*_max_, *V*_cmax_, and *J*_max_, were positively correlated with YOR (Fig. 5) and seed traits (Fig. 7). These results indicate that modern genotypes have greater biochemical capacity for carbon assimilation, irrespective of NPQ relaxation rates. A previous study of historical soybean lines did not consistently detect increases in *V*_cmax_, and *J*_max_ (Koester *et al*., 2016). However, these parameters represent potential capacity rather than fixed values, and their expression depends strongly on plant physiological status and environmental conditions (Farquhar, von Caemmerer and Berry, 1980; Walker *et al*., 2014). Importantly, our results show that photosynthetic capacity increased in modern genotypes despite unchanged NPQ relaxation. This pattern highlights a functional mismatch between biochemical capacity and photoprotective regulation: genotypes with greater potential for carbon fixation may not fully realize this capacity under natural field conditions, where irradiance fluctuates rapidly. Although photosynthesis was assessed here under steady-state conditions, previous studies have shown that slow NPQ relaxation can constrain carbon assimilation under fluctuating light by maintaining unnecessary energy dissipation (Kromdijk *et al*., 2016; De Souza *et al*., 2022; Beraldo, Alboresi and Morosinotto, 2026). Thus, the lack of improvement in NPQ relaxation suggests that part of the increased photosynthetic capacity in modern genotypes may remain underutilized in field conditions.

The mechanistic basis underlying differences between fast- and slow-relaxing genotypes cannot be explained by gene expression or pigment composition alone (Fig. 8). Although key NPQ components, including *VDE1*, *PsbS1*, and *ZEP1*, varied among genotypes, these differences did not match the relaxation dynamics used to classify fast- and slow-relaxing groups in the field. Gene expression and pigment analyses were performed at the R1/R2 stage after contrasting genotypes were identified based on NPQ relaxation at the V5/V6 stage, providing a mechanistic snapshot of these extreme phenotypes. Similarly, variation in pigment pools, including xanthophyll-cycle pool size and de-epoxidation state, did not explain differences in NPQ recovery. These results indicate that NPQ relaxation is governed primarily by dynamic regulatory processes rather than by the absolute abundance of molecular components. In particular, lumenal pH dynamics, enzyme activation kinetics, and protein-pigment interactions within the thylakoid membrane likely play central roles in controlling relaxation behavior (Murchie and Niyogi, 2011; Bassi and Dall’Osto, 2021). This may explain why genotypes with contrasting relaxation dynamics did not show consistent differences in gene expression or pigment composition.

Taken together, these findings identify NPQ relaxation as an underexplored dimension of photosynthetic performance in soybean. Breeding has enhanced carbon assimilation capacity and contributed to gains in seed number and seed weight, but it has not optimized the regulatory processes that determine how efficiently plants use this capacity under field conditions. Transgenic studies have demonstrated that accelerating NPQ induction and relaxation can increase carbon assimilation, biomass, and productivity under dynamic light environments (Kromdijk *et al*., 2016; De Souza *et al*., 2022), providing strong evidence that this limitation is biologically meaningful. Therefore, improving the speed and coordination of photoprotective recovery represents a promising strategy to better align regulatory flexibility with the high photosynthetic capacity of modern cultivars and enhance light-use efficiency and crop performance.

This study shows that soybean breeding increased photosynthetic capacity, reflected in higher *A*_max_, *V*_cmax_, and *J*_max_, and was associated with gains in seed number and seed weight. In contrast, NPQ relaxation showed no consistent directional change across genotypes released over the last century. This uncoupling indicates that gains in carbon assimilation were not matched by improvements in photoprotective regulation. As a result, modern genotypes may not fully use their enhanced photosynthetic capacity under field conditions. Although we examined multiple levels of organization, including gene expression, pigment composition, and physiological traits, the mechanisms underlying variation in NPQ relaxation remain unresolved. Dynamic processes such as lumenal pH regulation and xanthophyll-cycle kinetics likely play central roles but were not directly captured here. The substantial variation observed among genotypes highlights an opportunity to improve photoprotective regulation. Enhancing NPQ relaxation may help align regulatory flexibility with photosynthetic capacity and contribute to further gains in seed number and seed weight.

## Acknowledgment

We thank Desirae Aleman, Elena Pelech, Nichola Austen, Cibelle, Paulina Reynoso, Shuwan Xu, Mianchen Zhang, Miantong Zhang, and Mary Durstock for assisting with data collection, tissue sampling, and supporting in CO_2_ response curves. Bishal G. Tamang and Casey Kramer for LMA analysis. Jonathan Johnson for harvest procedure. Steven Burgess for VDE, ZEP, and PsbS primer design. Eliana Monteverde Dominguez for providing seeds.

## Author contributions

LPO, SPL, and EAA designed research; KA, LL, and LD quantified pigments and gene expression. LPO carried out the experiments, analyzed data, prepared figures and tables, and wrote the first draft; EAA, KA, LL and LD edited the draft; all authors read and approved the final version of the manuscript.

## Supplementary material

Supplementary material is available at Plant Physiology online

