## Supplementary material for "A century of soybean breeding increased photosynthetic capacity but not NPQ relaxation"

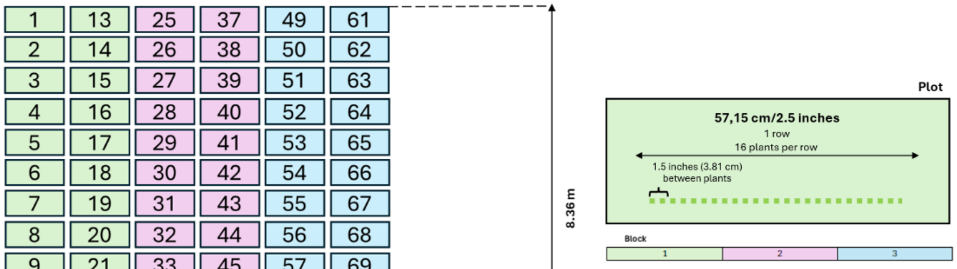


| | | | | | |
|---|---|---|---|---|---|
| Genotype1 | Genotype13 | Genotype12 | Genotype19 | Genotype22 | Genotype19 |
| Genotype2 | Genotype14 | Genotype11 | Genotype20 | Genotype23 | Genotype20 |
| Genotype3 | Genotype15 | Genotype10 | Genotype21 | Genotype24 | Genotype21 |
| Genotype4 | Genotype16 | Genotype9 | Genotype22 | Genotype1 | Genotype15 |
| Genotype5 | Genotype17 | Genotype8 | Genotype23 | Genotype2 | Genotype16 |
| Genotype6 | Genotype18 | Genotype7 | Genotype24 | Genotype3 | Genotype17 |
| Genotype7 | Genotype19 | Genotype6 | Genotype13 | Genotype4 | Genotype18 |
| Genotype8 | Genotype20 | Genotype5 | Genotype14 | Genotype5 | Genotype10 |
| Genotype9 | Genotype21 | Genotype4 | Genotype15 | Genotype6 | Genotype11 |
| Genotype10 | Genotype22 | Genotype3 | Genotype16 | Genotype7 | Genotype12 |
| Genotype11 | Genotype23 | Genotype2 | Genotype17 | Genotype8 | Genotype13 |
| Genotype12 | Genotype24 | Genotype1 | Genotype18 | Genotype9 | Genotype14 |

| Genotype ID | Genotype name |
|---|---|
| 1 | Dunfield |
| 2 | Illini |
| 3 | AK (Harrow) |
| 4 | Mandell |
| 5 | Adams |
| 6 | Ford |
| 7 | Shelby |
| 8 | Ross |
| 9 | Adelphia |
| 10 | Wayne |
| 11 | Calland |
| 12 | Williams |
| 13 | Woodworth |
| 14 | Zane |
| 15 | Thorne |
| 16 | Yale |
| 17 | Maverick |
| 18 | LD10 |
| 19 | LD11 |
| 20 | LD21-5547 |
| 21 | LD20-5413 |
| 22 | LD20-11665 |
| 23 | BC4_Narrow Leaf |
| 24 | BC4_Broad Leaf |

**Figure S1**. Field experimental layout used in this study. The experiment consisted of three blocks and a total of 72 plots. Plot size, row spacing, row length, and spacing between plots are indicated. **Alt text:** Diagram illustrating the experimental field layout and plant spacing used in the soybean trial. The left panel shows a numbered planting grid, while the upper right panel illustrates the spacing between plants within a row (1.5 inches or 3.81 cm) across a total row length of 57.15 cm (22.5 inches). The lower right panel presents the arrangement of soybean genotypes within the experimental layout, with repeated genotype labels distributed across planting positions.

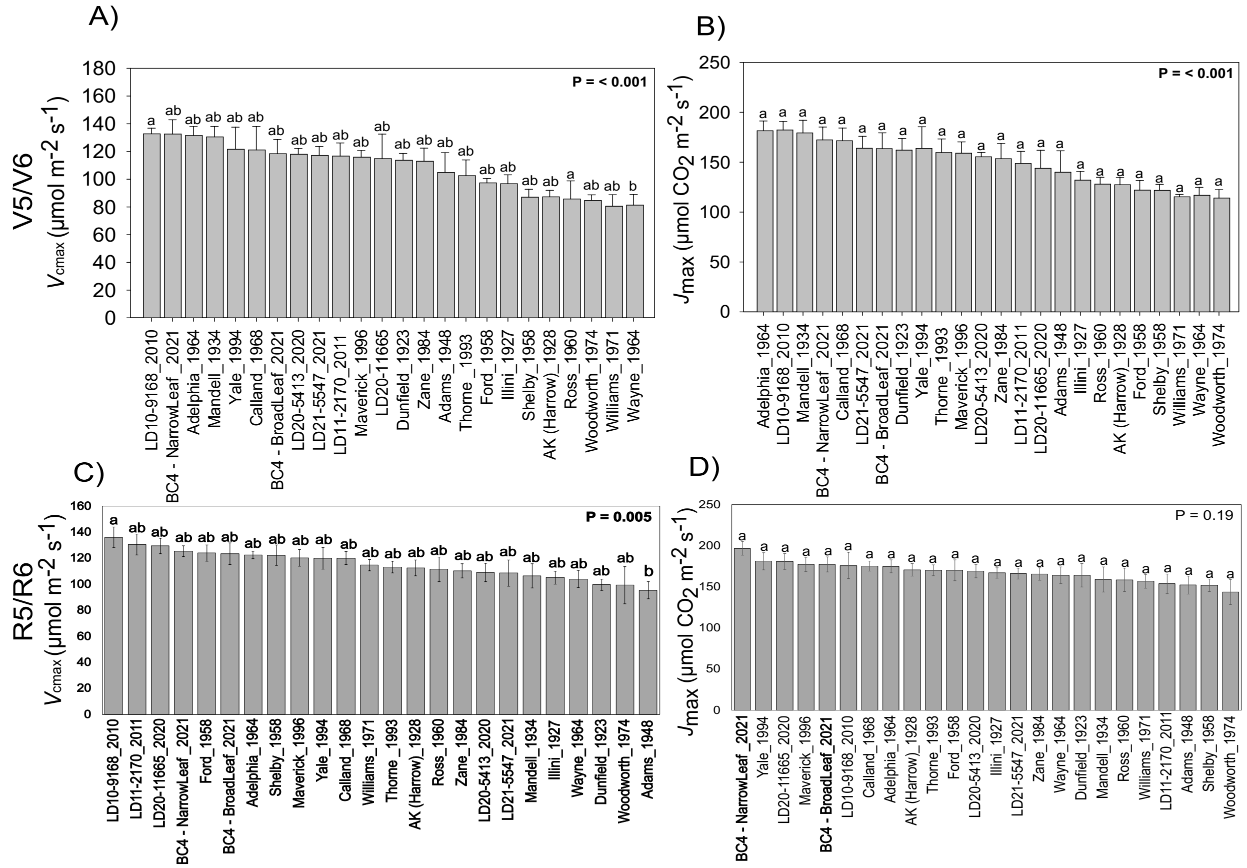


**Figure S2**. Genotypic variation in biochemical parameters of photosynthesis across 24 soybean genotypes. Maximum rate of carboxylation ($V_{cmax}$) and maximum rate of electron transport ($J_{max}$) were estimated from *A-Ci* curves at the vegetative (V5/V6) (A and B) and reproductive (R5/R6) stages (C and D). Genotypes are ordered by mean values within each panel. Data are means (± SE) of three replicate blocks ($n = 3$), each with two subsamples. Different letters indicate significant differences among genotypes (one-way ANOVA followed by Tukey's post hoc test, $P < 0.05$). The absence of letters in $J_{max}$ (R5/R6) indicates no significant genotypic effect ($P = 0.19$). Parameters were normalized to a leaf temperature of 25°C.

**Alt text:** Bar plots showing variation in photosynthetic biochemical parameters among 24 soybean genotypes at vegetative (top row) and reproductive (bottom row) developmental stages. Panels display maximum carboxylation rate ($V_{cmax}$) and maximum electron transport rate ($J_{max}$) estimated from A-Ci response curves and normalized to a leaf temperature of 25 °C. Genotypes are ordered according to mean parameter values within each panel. Bars represent mean values with error bars indicating standard error. Significant differences among genotypes are indicated by letters above bars, except for $J_{max}$ at the reproductive stage, where no significant genotypic effect was detected.

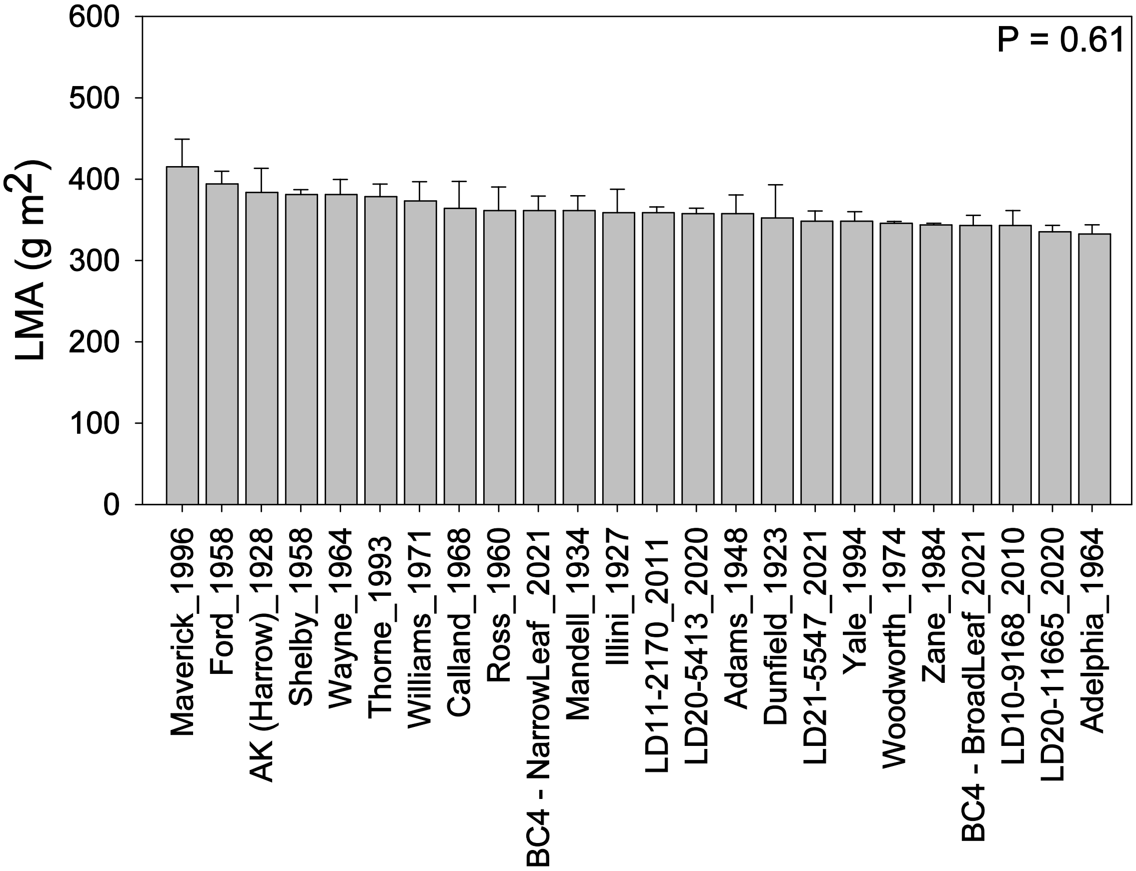


**Figure S3**. Leaf mass per area (LMA) across 24 soybean genotypes. LMA was calculated as the ratio of leaf dry mass to leaf area. Genotypes are ordered by mean values. Data are means (± SE) of three replicate blocks ($n = 3$), each with two subsamples. No significant differences were detected among genotypes (one-way ANOVA, P = 0.61).

**Alt text:** Bar plot showing leaf mass per area (LMA) across 24 soybean genotypes. Genotypes are ordered according to mean LMA values from highest to lowest. Bars represent mean values with error bars indicating standard error based on three replicate blocks with two subsamples each. LMA values were relatively similar among genotypes, and no significant differences were detected according to one-way ANOVA (P = 0.61).

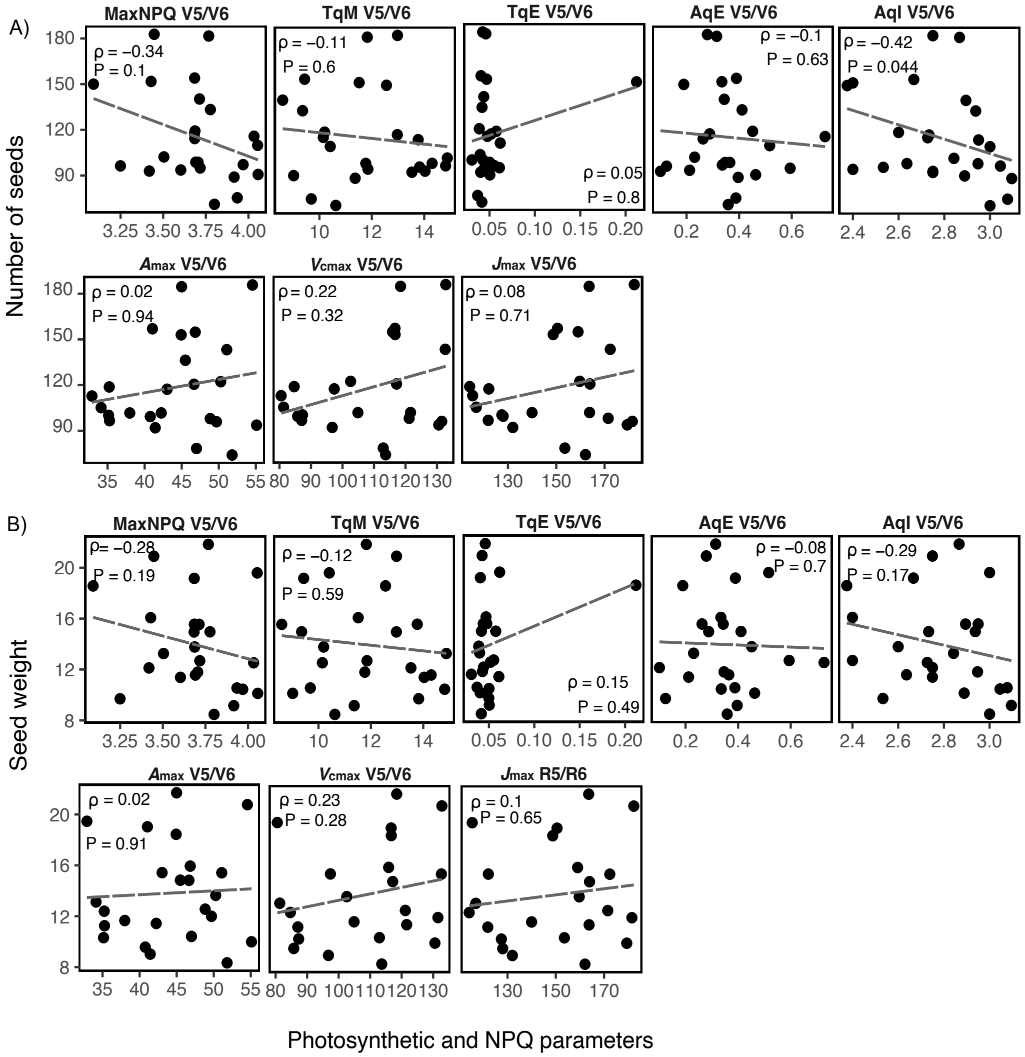


**Fig. S4**. **Scatter plots show relationships between NPQ parameters (Max NPQ, TqE, TqM, AqE, AqI), photosynthetic capacity ($A_{max}$, $J_{max}$ and $V_{cmax}$), and seed traits, seed number (A) and seed weight (B) per plant at the V5/V6**. Values represent the mean (± SE) of three replicate blocks (*n* = 3) with two subsamples. Lines show linear fits for visualization only; correlations were assessed using Spearman's rank correlation. Spearman correlation coefficients (ρ) and FDR-adjusted *P*-values (Benjamini–Hochberg correction, applied within each developmental stage) are shown in each panel. Dashed lines indicate non-significant correlations.

**Alt text:** Scatter plots showing relationships between seed traits and physiological parameters in soybean genotypes at the V5/V6 developmental stage. Panels compare seed number per plant (top two rows) and seed weight per plant (bottom two rows) with NPQ relaxation parameters, including Max NPQ, TqE, TqM, AqE, and AqI, as well as photosynthetic capacity parameters $A_{max}$, $V_{cmax}$, and $J_{max}$. Each point represents the mean value for an individual genotype, with dashed lines showing linear fits for visualization. Most relationships were weak and non-significant, although AqI showed a significant negative correlation with seed number. Spearman correlation coefficients and adjusted P-values are shown within each panel.

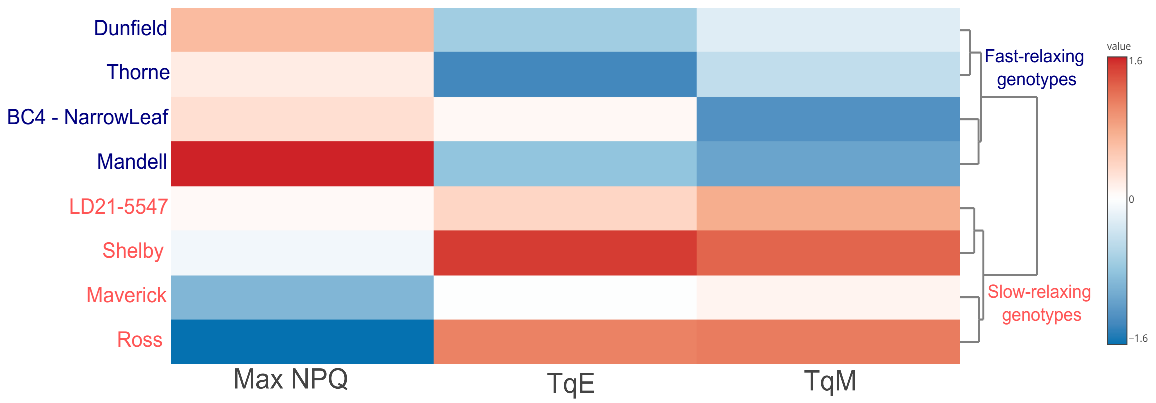


**Figure S5. Clustering of soybean genotypes based on NPQ relaxation kinetics highlights fast- and slow-relaxing groups.** This heatmap represents a subset of the full dataset shown in Figure 2, focusing on genotypes selected to emphasize contrasting extremes of NPQ relaxation components. Data are means (± SE) of three replicate blocks ($n = 3$), each with two subsamples at V5/V6 developmental stage. The heatmap shows variation in Max NPQ and relaxation kinetics (TqE and TqM) across selected soybean genotypes. Values were $\log_{10}$-transformed and autoscaled (Z-scores). Rows represent genotypes and columns represent NPQ parameters. The color scale indicates relative values (red = higher, blue = lower). Hierarchical clustering (Euclidean distance, Ward's method) separates genotypes into distinct groups characterized by fast NPQ relaxation (low TqE and TqM) and slow NPQ relaxation (high TqE and TqM). Genotype names are color-coded according to cluster assignment (blue = fast-relaxing genotypes; red = slow-relaxing genotypes).

**Alt text:** Heatmap showing normalized NPQ relaxation parameters among selected soybean genotypes classified as fast- or slow-relaxing types. Columns represent Max NPQ, TqE, and TqM, while rows represent individual soybean genotypes. Color intensity indicates relative parameter values, ranging from lower values in blue to higher values in red. Hierarchical clustering separates genotypes into groups with contrasting NPQ relaxation behavior, highlighting differences between fast- and slow-relaxing genotypes.

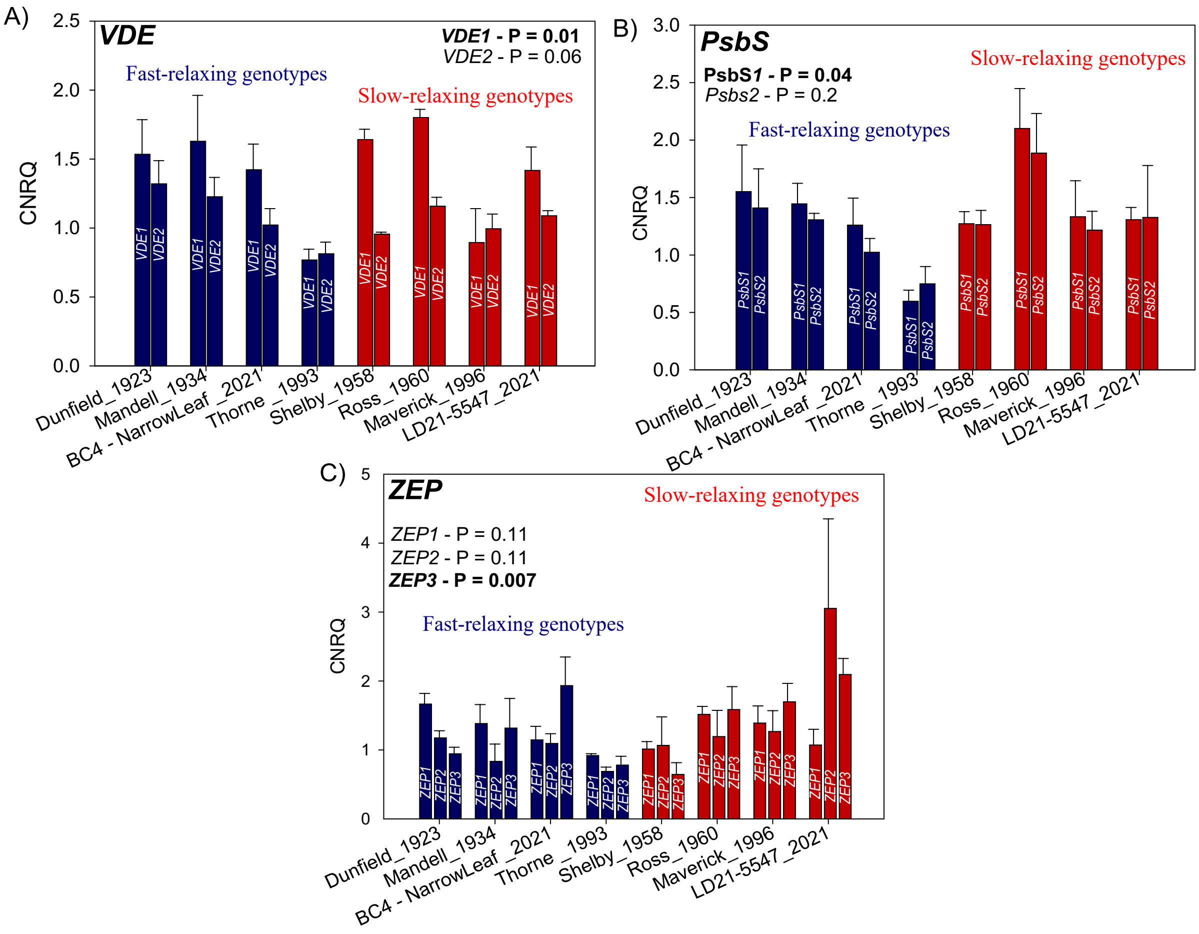


**Figure S6. Genotypic variation in expression of NPQ-related genes.** Transcript levels of *VDE1* and *VDE2* (A), *PsbS1* and *PsbS2* (B), and *ZEP1*, *ZEP2*, and *ZEP3* (C) were quantified in a representative subset of eight soybean genotypes contrasting in NPQ relaxation kinetics. Genotypes were previously classified as fast-relaxing (low TqE and TqM; blue bars) or slow-relaxing (high TqE and TqM; red bars). Data are presented as means (± SE) of three replicate blocks (*n* = 3), each with three subsamples. P-values from one-way ANOVA are indicated within each panel. Gene expression is reported as calibrated normalized relative quantity (CNRQ).

**Alt text:** Bar plots showing expression levels of photoprotective genes among soybean genotypes classified as fast- or slow-relaxing NPQ types. Panels display expression of *VDE* and *PsbS* gene family members across individual genotypes. Bars represent mean expression values with error bars indicating standard error. Slow-relaxing genotypes generally exhibited higher expression of *VDE* and *PsbS* genes compared with fast-relaxing genotypes. Statistical significance for total gene expression differences between groups is indicated by reported P-values.

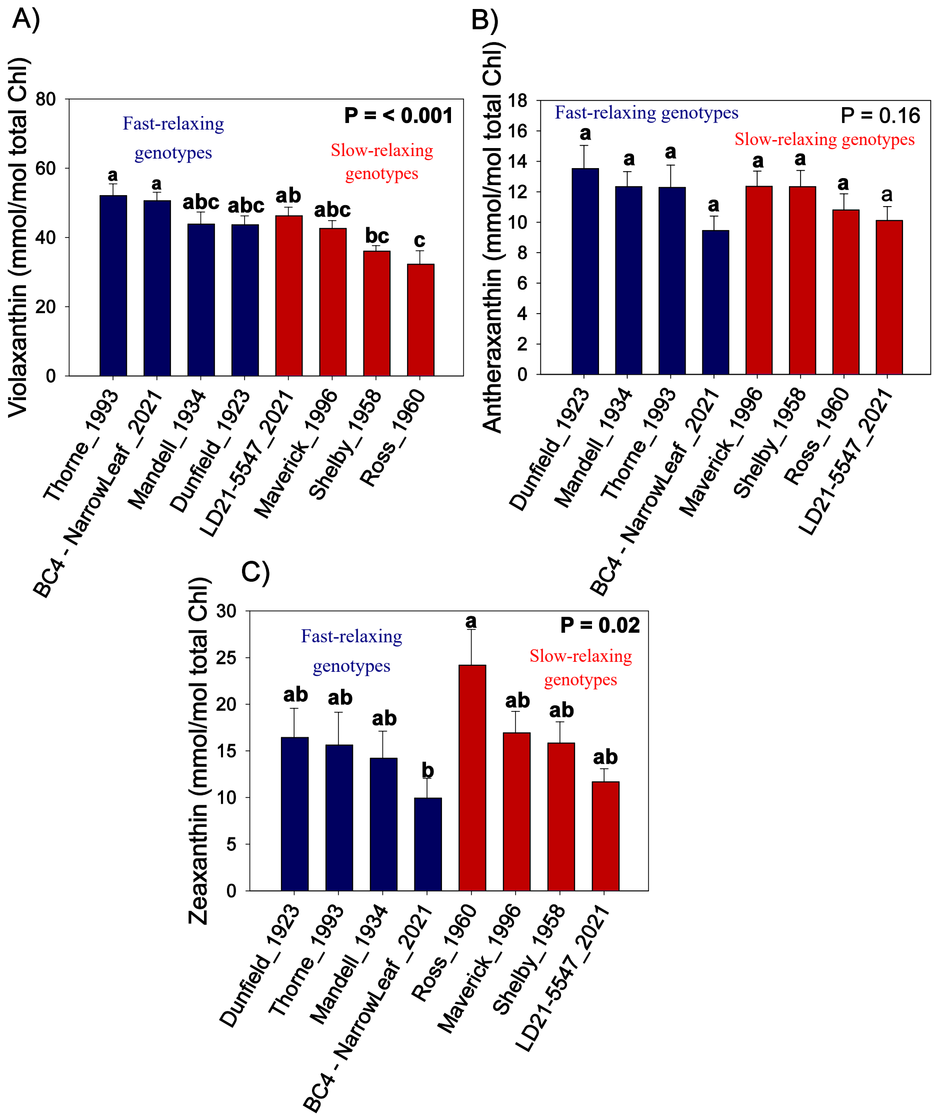


**Figure S7**. Genotypic variation in xanthophyll-cycle pigment composition. Violaxanthin, antheraxanthin, and zeaxanthin were quantified for a representative subset of eight soybean genotypes. Blue bars represent genotypes with historically higher Max NPQ and faster NPQ relaxation kinetics, while magenta bars represent genotypes with lower Max NPQ and slower relaxation kinetics. Data are presented as means (± SE) of three replicate blocks (*n* = 3), each with three subsamples. Significant differences (one-way ANOVA followed by posthoc Tukey's HSD test, $P < 0.05$) are indicated by letters. Pigment concentrations are expressed relative to total chlorophyll content (mmol/mol total Chl).

**Alt text:** Bar plots comparing xanthophyll-cycle pigment pools and de-epoxidation state between fast- and slow-relaxing soybean genotypes. Panels show measurements of total xanthophyll-cycle pigments and related pigment parameters across individual genotypes grouped by NPQ relaxation type. Bars represent mean values with error bars indicating standard error, and letters indicate significant differences among genotypes. Slow-relaxing genotypes generally exhibited higher pigment pool sizes and de-epoxidation levels than fast-relaxing genotypes.

**Table S1:** qPCR primers and assay conditions for reference genes and annotated *VDE*, *PsbS*, and *ZEP* paralogs in soybean (*Glycine max*; Gm).

| Gene ID [a] | Primer Name | Sequence (5' > 3') | Amplicon length (bp) | Taq Polymerase | Primer Conc. (nmol) | $T_{annealing}$ (°C) | Linear Range (ng)[b] | Efficiency (%) | Efficiency SE |
|---|---|---|---|---|---|---|---|---|---|
| Glyma.02G276600 | qPCR_GmELF1B_F | TTCACTCACTCACTCTGCACTCA | 91 | iTaq | 300 | 61.5 | 0.06 - 43.85 | 99.6 | 0.064 |
| | qPCR_GmELF1B_R | TGATGCCCTCTTCGGTGTGTA | | | 300 | | | | |
| Glyma.12G024700 | qPCR_GmCYP2_F | GCCGACTGTGGTCAACTCTC | 79 | iTaq | 300 | 61.5 | 0.02 - 43.85 | 104.1 | 0.083 |
| | qPCR_GmCYP2_R | GACACATTCAGAGCCACCGA | | | 300 | | | | |
| Glyma.03G253500 | qPCR_GmVDE1_F2 | TCCTCCCTAATCATTGCTTTTATGC | 127 | iTaq | 300 | 60 | 0.61 - 49.80 | 93.7 | 0.092 |
| | qPCR_GmVDE1_R2 | AGAGCACAGGCATCTAAACCT | | | 300 | | | | |
| Glyma.19G251000 | qPCR_GmVDE2_F4 | ACTGGCACTCCCAAAGTATGTG | 77 | iTaq | 300 | 60 | 0.08 - 49.80 | 94.5 | 0.055 |
| | qPCR_GmVDE2_R4 | AAACCTCTAGTCCTGTGAAACCTTA | | | 300 | | | | |
| Glyma.17G174500 | qPCR_GmZEP1_F2 | TTCTCACACAAACTGCAACCAT | 140 | iTaq | 300 | 60 | 0.02 - 45.01 | 97.4 | 0.074 |
| | qPCR_GmZEP1_R2 | GCCAACAACAAAAGGTGAAGCA | | | 300 | | | | |
| Glyma.11G055700 | qPCR_GmZEP2_1F | AGGTGACAAGATCTGCTTCAAGT | 142 | SSO | 500 | 57 | 0.35 - 11.2 | 103.0 | 0.082 |
| | qPCR_GmZEP2_1R | GCTCTACGGCTCCGACTTTT | | | 500 | | | | |
| Glyma.09G000600 | qPCR_GmZEP3_F2 | TTGGGTCAGCCAAATTCCAA | 84 | iTaq | 300 | 60 | 0.14 - 45.01 | 97.8 | 0.140 |
| | qPCR_GmZEP3_R2 | GCACAGAGGATTTGCACAGAA | | | 300 | | | | |
| Glyma.06G113200 | qPCR_GmPsbS2_F3 | TGCATTGAGTTCGTCGTAAACA | 75 | iTaq | 300 | 60 | 0.02 - 51.87 | 96.1 | 0.056 |
| | qPCR_GmPsbS2_R3 | GTCTAATCATGACATCGAGACGC | | | 300 | | | | |
| Glyma.04G249700 | qPCR_GmPsbS1_F6 | CGTTTGCTCATCTATCATAAAATGC | 140 | iTaq | 500 | 57 | 0.05 - 100.83 | 92.6 | 0.055 |
| | qPCR_GmPsbS1_R6 | TGGCAAATCGAAGCACTATGA | | | 500 | | | | |

[a] Phytozome v13 *Glycine max* Wm82.a2.v1 gene identifier (2020-08-24)

[b]Linear Range is representative of amount of RNA added to qPCR reaction assuming a 100% reverse transcription efficiency during cDNA synthesis.

**Table S2: qPCR Primer Characteristics**

| Primer Name | BLAST alignments[a] | Spliceforms[b] | Primer Location | Location[c] | $T_m$ °C [d] | ΔG homodimer[e] | ΔG heterodimer[f] | Tm hairpin [g] | Mfold / Tm °C [h] | Source |
|---|---|---|---|---|---|---|---|---|---|---|
| qPCR_GmELF1B_F | NM_001249608.2 | 1 | Exon 1 | Exon 1 | 65.5 | -7.05 | -5.09 | 6.5 | 47.1 | doi: 10.1126/science.adc9831 |
| qPCR_GmELF1B_R | | | Exon 1 | | 65.6 | -3.61 | | 32 | | |
| qPCR_GmCYP2_F | NM_001357079.1 | 1 | Exon 1 | Exon 1 | 64.6 | -3.61 | -3.67 | 44.8 | 41.7 | doi: 10.1126/science.adc9831 |
| qPCR_GmCYP2_R | | | Exon 1 | | 64.3 | -3.61 | | 15.7 | | |
| qPCR_GmVDE1_F2 | Not Available | 5 | 3'UTR | 3'UTR | 62.5 | -3.43 | -6.69 | 29.4 | 54 | This publication |
| qPCR_GmVDE1_R2 | | | 3'UTR | | 59.5 | -4.67 | | 47.1 | | |
| qPCR_GmVDE2_F4 | XM_014771336.2, NM_001254020.1 | 1 | Exon 2 | Exon 2 | 62.1 | -5.02 | -3.54 | 40.5 | 41.1 | This publication |
| qPCR_GmVDE2_R4 | | | Exon 2 | | 62.5 | -4.16 | | 26.3 | | |
| qPCR_GmZEP1_F2 | Not Available | 1 | 5' UTR | Exon 1 | 58.4 | -7.05 | -7.05 | 21.9 | 55.9 | This publication |
| qPCR_GmZEP1_R2 | | | Exon 1 | | 60.1 | -7.05 | | 40.2 | | |
| qPCR_GmZEP2_1F | Not Available | 1 | Exon 16 | Exon 16 | 60.9 | -7.82 | -4.88 | 32 | 55.9 | This publication |
| qPCR_GmZEP2_1R | | | 3'UTR | | 60.5 | -6.68 | | 52.2 | | |
| qPCR_GmZEP3_F2 | XM_006586667.3, XM_003534289.4 | 6 | 5' UTR | Exon 1 | 56.4 | -6.97 | -7.32 | 47.7 | 57.7 | This publication |
| qPCR_GmZEP3_R2 | | | Exon 1 | | 59.5 | -7.05 | | 32.9 | | |
| qPCR_GmPsbS2_F3 | Not Available | 1 | Exon 4 | Exon 4 | 58.4 | -7.05 | -6.53 | 26.6 | 52.3 | This publication |
| qPCR_GmPsbS2_R3 | | | 3'UTR | | 62.9 | -8.53 | | 29.4 | | |
| qPCR_GmPsbS1_F6 | Not Available | 1 | Exon 4 | Exon 4 | 60.9 | -3.89 | -8.98 | 21.3 | 53.7 | This publication |
| qPCR_GmPsbS1_R6 | | | 3'UTR | | 57.5 | -6.76 | | 32.1 | | |

[a] Primer-BLAST (https://www.ncbi.nlm.nih.gov/tools/primer-blast/index.cgi) alignment. Searching Refseq RNA database, species Glycine max (taxid:3847)

[b] Number of splice forms based on Phytozome v13 *Glycine max* Wm82.a2.v1 gene identifier (2020-08-24). .

[c] Location of amplicon

[d] $T_m$ estimated using IDT OligoAnalyzer (https://www.idtdna.com/calc/analyzer/) conditions ([Oligo] 0.33 µM, 50 mM $Na^{2+}$, 3 mM $Mg^{2+}$, 1.2 mM dNTPs).

[e,f,g,] Estimated using IDT OligoAnlyzer (https://www.idtdna.com/calc/analyzer/) conditions (Oligo 0.3uM, 50 mM Na2+, 3 mM Mg2+, 0.8 dNTPs).

[g]The highest Tm is reported for potential hairpin structures.

[h] Mfold analysis of DNA amplicon structure. The highest Tm is reported for potential secondary structures.
